# Benchmarking circRNA Detection Tools from Long-Read Sequencing Using Data-Driven and Flexible Simulation Framework

**DOI:** 10.1101/2025.04.17.649290

**Authors:** Anastasia Rusakovich, Sebastien Corre, Edouard Cadieu, Rose-Marie Fraboulet, Victor Le Bars, Marie-Dominique Galibert, Thomas Derrien, Yuna Blum

## Abstract

Circular RNAs (circRNAs) are unique non-coding RNAs with covalently closed loop structures formed through backsplicing events. Their stability, tissue-specific expression patterns, and potential as disease biomarkers have garnered increasing attention. However, their circular structure and diverse size range pose challenges for conventional sequencing technologies. Long-read Oxford Nanopore (ONT) sequencing offers promising capabilities for capturing entire circRNA molecules without fragmentation, yet the effectiveness of bioinformatic tools for analyzing this data remains understudied.

This study presents the first comprehensive benchmark comparison of three specialized tools for circRNA detection from ONT long-read data: CIRI-long (Zhang et al., 2021), IsoCIRC (Xin et al., 2021), and circNICK-Irs (Rahimi et al., 2021). To address the lack of standardized evaluation frameworks, we developed a novel computational pipeline, open-source and freely available, to generate realistic simulated circRNA ONT long-read datasets. Our pipeline integrates several molecular features of circRNAs extracted from established databases and real datasets into NanoSim tool (Hafezqorani et al., 2020) and outputs FASTQ reads reflecting therefore biological diversity and technical properties.

We systematically assessed tool performance across key metrics, including precision, recall, specificity, accuracy, and F1 score. Our analysis revealed distinct performance profiles: while all tools exhibited high specificity, they varied in precision and their ability to detect different circRNA subtypes, often showing limited sensitivity and precision. Notably, the overlap in detected circRNAs among tools was relatively low. Additionally, computational efficiency varied significantly across the tools. This suggests that relying on a single tool might not be ideal, and combining tools or improving algorithms could be necessary for more accurate circRNA detection from ONT data.

This benchmark provides valuable insights for researchers selecting appropriate tools for circRNA studies using ONT sequencing. Furthermore, our customizable simulation framework, offering a resource to optimize detection approaches and advance bioinformatic tool development for circRNA research is freely available at: https://gitlab.com/bioinfog/circall/nano-circ.

## Introduction

Non-coding RNAs (ncRNAs), which make up a significant portion of the transcriptome, have emerged as critical players in various cellular processes, ranging from gene regulation to disease pathogenesis (Esteller, 2011). Among them, circular RNAs (circRNAs) represent a unique class of ncRNAs, formed through non-canonical backsplicing events, resulting in a covalently closed loop structure. Since their discovery, circRNAs have garnered increasing attention due to their stability, tissue-specific expression, and potential roles as biomarkers in diseases, including cancer (Anastasiadou et al., 2018; Verduci et al., 2021). However, their circular structure and diverse size range - spanning from less than 100 to almost 100000 nucleotides (Rybak-Wolf et al., 2015) - pose significant challenges to conventional sequencing technologies, particularly second-generation sequencing methods, which often rely on read fragmentation.

Recent advancements in third-generation sequencing technologies, such as Oxford Nanopore sequencing, have provided novel opportunities to explore circRNAs in greater detail (Wang et al., 2021). Unlike second-generation methods, Nanopore sequencing offers long-read capabilities that can capture entire circRNA molecules without the need for fragmentation, making it a promising approach for the comprehensive characterization of circRNAs. However, the full potential of long-read sequencing for circRNA discovery and annotation depends on an integration of wet-lab approach, sequencing platform and bioinformatics tools that can accurately detect and quantify circRNAs from these datasets.

Despite the growing number of bioinformatics tools developed for circRNA detection (Drula et al., 2024), a comprehensive benchmark comparing their performance on long-read Nanopore sequencing data remains absent. To address this gap, we performed a systematic comparison of 3 circRNA detection tools - CIRI-long (Zhang et al., 2021), IsoCIRC (Xin et al., 2021) and circNICK-lrs (Rahimi et al., 2021).

We selected these tools because they represent the three major methodological approaches to long-read circRNA detection, each with distinct experimental protocols and computational pipelines: rolling circle reverse transcription (CIRI-long), rolling circle amplification followed by nanopore sequencing (isoCirc), and direct circRNA linearization approaches (circNick-LRS). This selection provides comprehensive coverage of the current landscape of long-read circRNA detection methodologies, enabling evaluation of how different experimental and computational strategies affect detection performance across various circRNA characteristics.

The complexity of circRNA detection requires a reproducible evaluation framework that goes beyond traditional wet-lab datasets. Such datasets, experimentally generated from biological samples using standard circRNA enrichment and sequencing protocols, inherently lack an established ground truth and suffer from limitations such as biological variability and sequencing biases. Simulated datasets offer a solution to these challenges, providing a controlled environment with precisely known circRNA and linear RNA annotations. By generating in silico reads that mimic the molecular and sequencing characteristics of Nanopore technologies, we can create a comprehensive ground truth dataset that allows for rigorous assessment of performance by each of circRNA detection tools.

In this study, we provide a first framework for the generation of simulation data of long-read sequencing of circRNA and present the benchmark analysis of these 3 tools using simulated datasets. By evaluating their performance across different metrics, we aim to provide valuable insights into the strengths and limitations of each tool, guiding researchers in selecting the most appropriate software for their circRNA studies (Fig.1).

**Figure 1.**
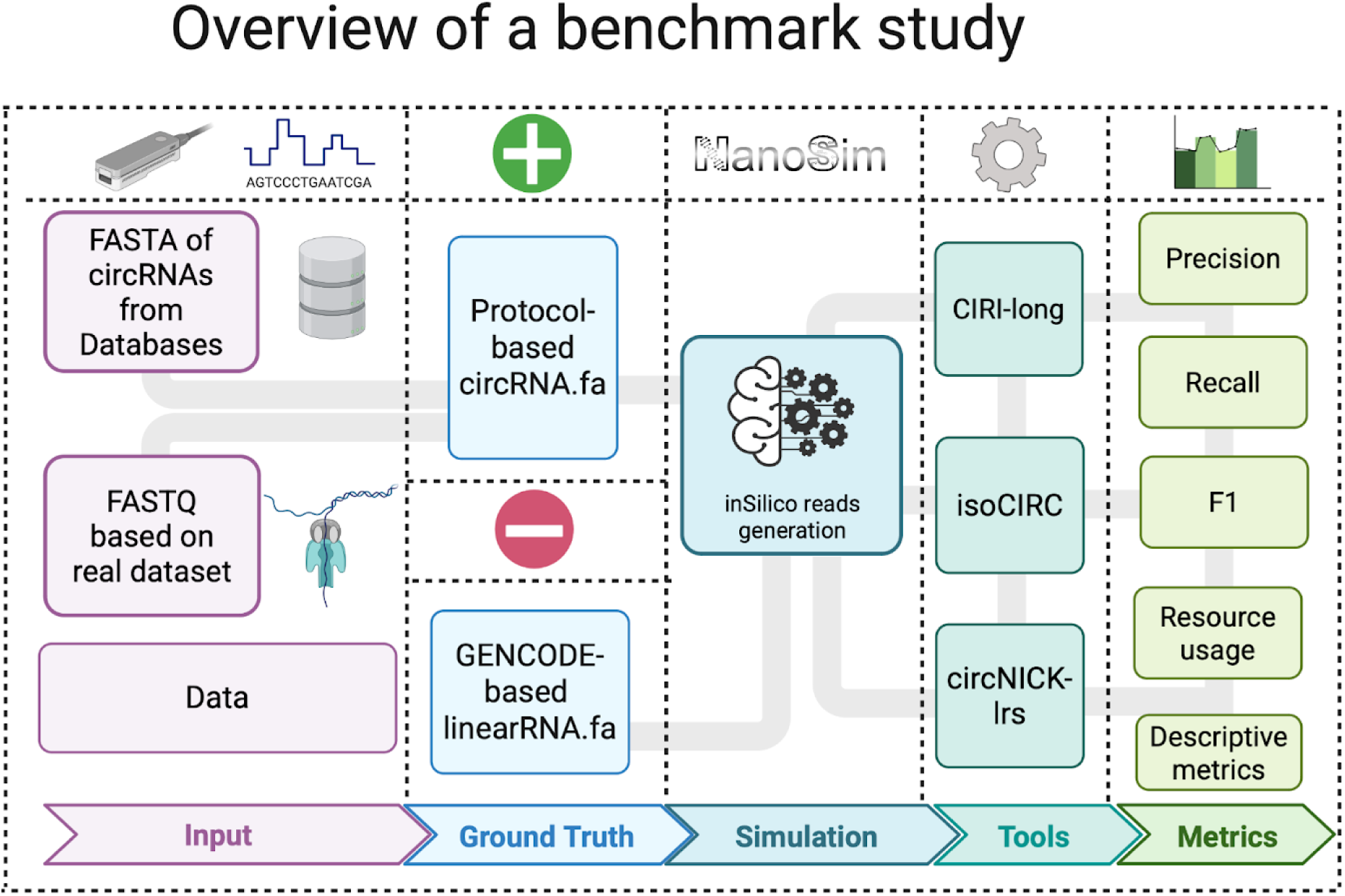
Overview of the benchmarking study. Schematic representation of the benchmarking framework for circular RNA (circRNA) detection tools. The workflow progresses through five stages: (I) input FASTQ sequences from circRNA databases (see Feature extraction from databases section) and real wet lab protocols (see Mouse brain dataset section) that are used for feature extraction and preparation of ground truth; (II) ground truth preparation using custom scripts for positive ground truth and GENCODE based linear RNA sequences as negative ground truth; (III) in silico FASTQ read simulation with NanoSim tool; (IV) computational analysis using CIRI-long, isoCirc, and circNICK-lrs tools; (V) performance evaluation through multiple metrics including sensitivity, precision and F1 score, calculated across different overlap thresholds. Figure created in https://BioRender.com.

## Methods

### Wet lab protocols for circular RNA identification using long-read sequencing data

The three wet lab protocols generate distinct read types with specific bioinformatic requirements (see Supplementary Materials). CIRI-long (rolling circle reverse transcription) and isoCirc (rolling circle amplification) both produce concatemeric reads with multiple circRNA copies per read; their pipelines detect tandem repeat patterns and generate consensus sequences. circNICK-LRS produces single-copy linearized reads spanning back-splice junctions and uses split-read alignment for BSJ detection (Supplementary Fig.2). Size selection ranges differ across protocols: CIRI-long targets ∼1 kb fragments (AMPure XP beads), isoCirc selects 3–50 kb fragments (BluePippin), and circNICK-LRS selects 350 bp–10 kb fragments (agarose gel electrophoresis).

**Figure 2.**
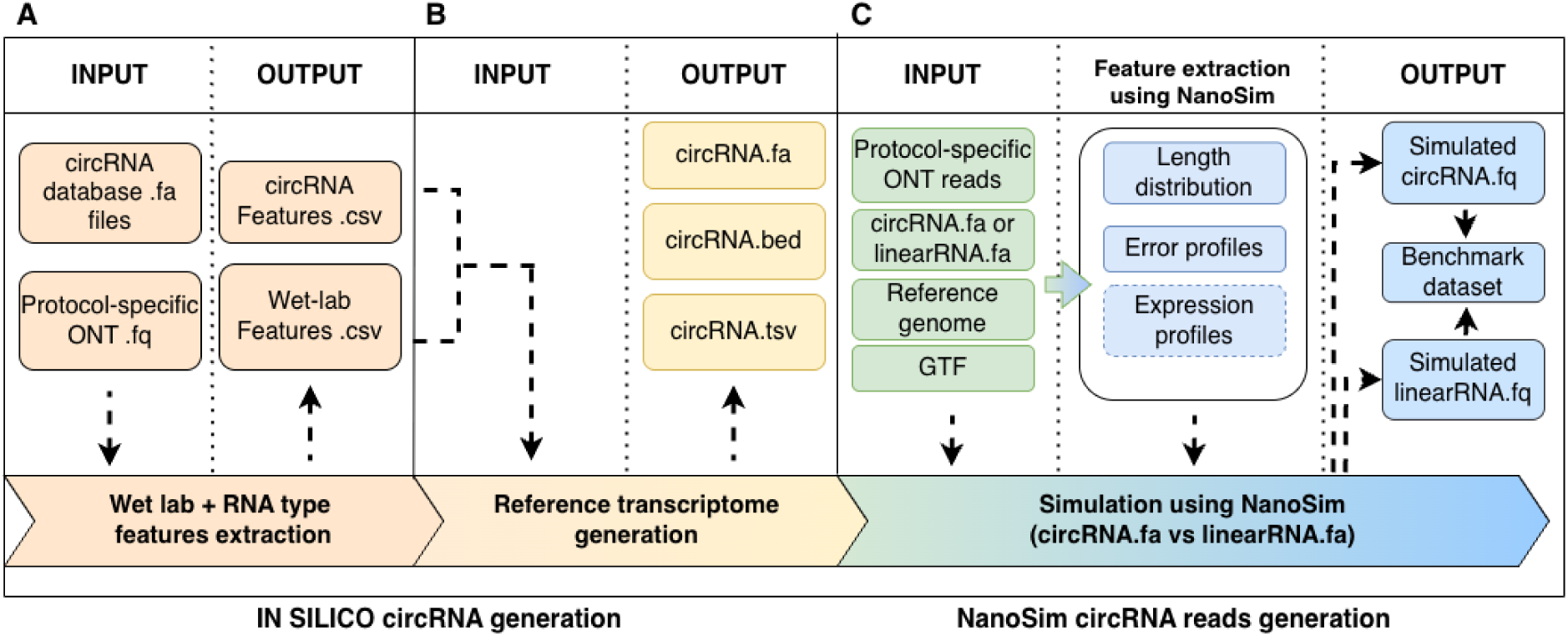
Computational Workflow for Nanopore Sequencing circRNA Simulation. The workflow integrates wet-lab Oxford Nanopore Technologies (ONT) read characteristics with computational simulation techniques. Panels A-C depict sequential stages of data processing and simulation: (A) initial feature extraction from circRNA databases (more in circRNA databases section) and experimental reads (more in wet lab dataset section), (B) generation of protocol-specific reference circRNA sequences, and (C) NanoSim-based simulation generating *in silico* reads for circRNA and linear RNA that are subsequently pooled into benchmark dataset.

### Tools for circular RNA identification using long-read sequencing data

**CIRI-long (Zhang et al., 2021)**: is a computational method designed for profiling circRNAs using nanopore long-read sequencing data based on a rolling circle reverse transcription approach. The algorithm in this study reconstructs full-length circRNA sequences through a multi-step process: (1) Data preprocessing: demultiplexing, quality control, and adapter removal using guppy (v3.3.0), Porechop (v2.0.4), and pycoQC (v2.5.0.14); (2) Repetitive pattern identification: k-mer-based detection of circular patterns using k=8 and k=11 with homopolymer-compressed k-mers; (3) Consensus sequence generation: partial order alignment (SPOA) to create cyclic consensus sequences with 80% similarity threshold between repetitive segments; (4) Mapping and BSJ detection: alignment using mappy (v2.17) for sequences >150 bp and bwapy (v0.1.4) for shorter sequences, followed by iterative alignment strategy; (5) Filtering and validation: canonical splice signal validation (GT/AG, GC/AG, AT/AC), an 80% sequence similarity threshold and clustering based on genomic coordinates. The method validates circRNAs against the circAtlas database or other user-provided databases. More details can be found at [https://github.com/bioinfo-biols/CIRI-long].

**isoCirc (Xin et al., 2021)**: is a method for sequencing and characterizing full-length circular RNA isoforms using rolling circle amplification followed by nanopore long-read sequencing. The computational pipeline in this study involves: (1) Data preprocessing: demultiplexing, quality control, and adapter removal using guppy (v2.1.3); (2) Repetitive pattern identification: Tandem Repeat Finder (v4.0.9) to identify multiple copies of circRNA sequences within reads; (3) Consensus sequence generation: construction of consensus sequences from detected tandem repeats with copy number-dependent error correction (average copy number of 14.5); (4) Mapping and BSJ detection: alignment of consensus sequences to reference genome using minimap2 (v2.17), with BSJ identification through split-read analysis; (5) Filtering and validation: multi-tiered alignment scoring, stringent validation of back-spliced junctions (BSJs) and forward-spliced junctions (FSJs), requiring high mapping quality and fidelity. The method characterizes alternative splicing events and validates against circBase and MiOncoCirc databases. More information can be found at [https://github.com/Xinglab/isoCirc].

**circNick-LRS (Rahimi et al., 2021)**: is a computational workflow for profiling circRNAs from linearized circRNA nanopore long-read sequencing data. The pipeline in this study consists of: (1) Data preprocessing: demultiplexing, quality control and length filtering using Guppy (v3.4.5) and NanoFilt (v2.6.0); (2) Mapping: direct alignment to reference genome using pblat against the human (hg19) or mouse (mm10) genome; (3) BSJ detection: identification of back-splice junctions utilizes pblat (v35) and bedtools (v2.29.2), requiring a minimum Blat score of 30 on both sides of the BSJ; (4) Filtering and validation: retention of reads mapping to same strand, within 1 Mb distance, non-overlapping by ≥50 bp, and in reverse genomic order; (5) Annotation and classification: aligning reads to the closest refSeq exon (within 30 bp), and validation against multiple databases while reads overlapping annotated circRNAs (≥95%) are corrected and retained. The method validates against circBase, circAtlas, and CIRCpedia databases. Further details can be found at [https://github.com/omiics-dk/long_read_circRNA].

More information about these tools is available in Supplementary table 1.

### Wet-lab dataset

#### Mouse brain dataset

We selected the CIRI-long protocol and mouse brain dataset from CIRI-long study as our basis to establish a robust foundation for our benchmarking study. The CIRI-long paper has the highest number of citations and provides the most comprehensively described wet-lab conditions, making it an ideal reference point for comparative analysis. Additionally, mouse circRNA databases contain over one million well-curated annotations, providing robust ground truth data essential for reliable feature extraction and validation of detection accuracy across all three methods.

The FASTQ file from the Zhang et al. study (2021) was obtained from the National Genomics Data Center (China National Center for Bioinformation) under the accession number CRA003317 [https://ngdc.cncb.ac.cn/gsa/browse/CRA003317]. This dataset, containing 2,746,616 reads and sequencing quality Q20, served as the foundation for our simulation parameters and error modeling.

### Ground truth construction

#### circRNA databases

We used two complementary circRNA databases: circAtlas v.3 (Wu et al., 2020) and circBase (Glažar et al., 2014) (more information about databases is available in Supplementary materials) based on four key criteria: (1) Multi-organism support - both databases provide mouse and human circRNA annotations; (2) Condition-independent curation - entries represent general circRNA populations rather than condition-specific datasets; (3) Sequence availability - both provide mature circRNA sequences in FASTA format that allowed us to extract features from them; and (4) Comprehensive coverage - high numbers of entries (circBase: >140,000; circAtlas: >3 million) ensure robust statistical analysis. Importantly, we used the intersection of these databases to enhance confidence in our reference dataset, as circRNAs supported by both independent databases are less likely to represent study-specific artifacts and more likely to constitute authentic circRNA sequences.

#### Genome assembly

We utilized the GRCm38.p4 mouse genome assembly, since circNICK-lrs tool is compatible only with this version of the reference genome. Reference genome sequences and genomic annotations were obtained from the GENCODE consortium (Mudge et al., 2025), specifically using the GENCODE version M10 mouse gene annotation file. This version provides comprehensive gene models including protein-coding genes, long non-coding RNAs, and pseudogenes, ensuring complete coverage for circRNA classification and validation across all genomic contexts (exonic, intronic, and intergenic regions).

### Data simulation

#### Feature extraction from databases

To understand the composition of circular RNAs, we have analysed mouse circRNAs found at an intersection between circRNA databases. We developed a computational pipeline to extract features from circRNA annotations using Python-based bioinformatics tools (pybedtools, pysam, pandas). The analysis integrated genomic coordinates, sequence information, and transcriptomic context from input BED files, reference genomes (FA), and gene annotation (GTF) files (Fig. 2A). Our methodology extracted key circRNA characteristics, including genomic location (chromosome, start, end, strand), mature RNA length, and splice site details. We classified splice sites based on canonical motifs (GT-AG, GC-AG, AT-AC) and performed intersectional analysis to annotate gene and transcript types.

#### Feature extraction from nanopore sequences

To understand the effect of wet lab protocol on sequencing data, we developed a Python-based computational pipeline to extract and visualize detailed sequencing characteristics from FASTQ files (Fig. 2A). The analysis quantified read metrics including length distributions and repeat patterns present in the wet lab data. Data processing and statistical analysis were performed using Python libraries including pandas and NumPy for handling structured datasets and numerical operations, while visualization utilized Python plotting libraries matplotlib and seaborn for generating distribution plots and statistical comparisons. Rolling circle amplification characteristics were analyzed using Tandem Repeats Finder (TRF v4.09) integrated within our Python framework to detect rolling circle periods, estimate copy numbers and identify repeat patterns.

#### in-Silico circRNA generation

Building upon our database and wet-lab protocol feature extraction pipelines, we developed an approach to simulate circular RNA types (Fig. 2B). Using genomic features extracted from exonic, intronic, and intergenic regions, we generated four circRNA types: exonic circRNAs (ecircRNAs), circular intronic RNAs (ciRNAs), exon-intron circRNAs (EIciRNAs) and intergenic circRNAs (Supplementary fig.1). Key generation features included random sequence extraction, length-constrained generation, splice site preference, type specific randomisation of exon skipping and intron retention events and rolling circle amplification.

To capture the biological complexity of circRNA isoforms, our simulator incorporates alternative splicing patterns that reflect natural circRNA diversity. For eciRNAs, we implemented exon skipping events with 10% probability for each exon, allowing generation of multiple isoforms from the same genomic locus with varying exonic compositions. EIciRNAs exhibit more complex splicing patterns with 15% probability of exon skipping and 70% probability of intron retention for each intron, reflecting the characteristic exon-intron structure that defines this circRNA class. ciRNAs are generated from intronic sequences with predominantly single-exon structures (59.4%) representing complete intron circularization, while multi-exonic ciRNAs result from combining multiple intronic regions within the same gene. Intergenic circRNAs are generated from non-genic genomic regions with random positioning and exhibit variable exon counts (35% single-exon, 65% multi-exonic) based on the extracted distribution patterns from our database analysis. These type-specific structural probabilities were derived from our database analysis and ensure that simulated circRNAs exhibit the structural diversity observed in real biological systems, including the presence of multiple isoforms per locus that challenge detection tools differently.

We implemented sequence quality filtering to exclude sequences with more than 10% unidentified nucleotides (N bases or gap in the assembly). The generation process dynamically selected sequence start positions, controlled rolling circle replication defined by user parameters, and captured detailed metadata including transcript identifiers, gene coordinates, and strand information. To validate the simulated circRNAs, we employed BLAT [14] for quality control validation of backsplice junctions. While BLAT is less optimal for long-read alignment compared to minimap2, we selected it for this validation step because its output format displays clearly in genomic browsers (e.g. UCSC [15], IGV [16]), enabling straightforward manual validation of back-splice junctions and visual identification of the characteristic “tail-before-head” pattern of circRNA reads during quality control. It is important to note that BLAT served solely as a quality control validation tool for our simulated data and was not used for precise genomic mapping in the benchmarking analysis, where minimap2 was employed for its superior optimization with nanopore data. This validation step ensures that our simulated circRNAs maintain the structural characteristics necessary for downstream tool evaluation while incorporating the isoform complexity that distinguishes transcriptome-level detection (general boundary identification) from full-length isoform detection (complete sequence reconstruction with accurate internal structure).

More information about the used features and output file can be found in Supplementary table 2 and 3.

#### in-silico circRNA long-reads data simulation

We utilized NanoSim version 3.2.3 to simulate Nanopore sequencing reads from the circRNA sequences generated in our previous simulation step to use as positive ground truth and from linear RNA sequences to use as negative ground truth (Fig.2C). The simulation uses multiple input files: a control FASTQ file (CRR194180.fq), reference genome (GRCm38.p4), and transcriptome annotation (gencode.vM10). The NanoSim simulation process involved four stages: read analysis for characterizing sequencing properties using the control FASTQ file with minimap2 alignment, circRNA read simulation generating 200,000 reads from our previously created circRNA.fa file, transcriptome quantification for analyzing expression levels, and linear read simulation generating 200,000 linear RNA reads from transcriptome annotation.

We selected the Guppy basecaller option because it represents the most accurate basecaller for ONT data, ensuring our simulated reads reflect contemporary sequencing quality compared to the available older alternative (Albacore). The cDNA 1D read type was chosen to match the library preparation method used by all benchmarked tools and represents the current ONT standard. We chose minimap2 for alignment during characterization because it provides superior accuracy for long-read transcriptome data compared to alternative LAST aligner. Importantly, we disabled NanoSim’s intron retention modeling to prevent conflicts with our custom circRNA-specific simulation scripts, allowing us to control splicing patterns precisely according to our circRNA type-specific generation rather than relying on generic intron retention models.

NanoSim learns error profiles (insertion, deletion, and substitution rates) from the real ONT sequencing data provided during the training phase, rather than applying fixed error rates. Error profiles were derived from the Zhang et al. (2021) mouse brain FASTQ file (CRR194180.fq), resulting in simulated reads with average quality scores of approximately Q20 and a chimeric read proportion consistent with the real dataset. Output simulated reads are in FASTQ format and untrimmed. (Fig. 2C)

### Benchmark Datasets

The benchmark datasets were generated for *Mus musculus* and used 7,503 unique simulated circRNAs across four types (eciRNA, EIciRNA, ciRNA, intergenic) and 117,667 unique linear transcripts (GENCODE vM10). Sequencing reads were simulated using NanoSim with the following parameters: ONT platform, cDNA 1D library preparation, Guppy basecaller. Q-score ∼20, error profiles and homopolymer modeling were derived from the Zhang et al. (2021) mouse brain dataset.

Four datasets were generated to assess performance across varying conditions. Three balanced datasets contained equal proportions of circRNA and linear RNA reads at increasing sequencing depths: 100,000/100,000 reads (low depth, presented in Supplementary materials with a few reads icon), 200,000/200,000 reads (medium depth, referred to as 50/50 mode, presented in the main manuscript with a 50/50 icon), and 300,000/300,000 reads (high depth, presented in Supplementary materials with a block of many reads icon). One realistic dataset mimicked circRNA enrichment conditions with approximately 3% circRNA reads (12,000 circRNA / 380,000 linear RNA reads), referred to as realistic mode, presented in the main manuscript with a pipette icon. Each dataset includes three independent simulation replicates to account for stochastic variability in read generation and tool execution.

### circRNA detection tools parameters

For the circRNA detection step we used the following versions of the tools: CIRI-long v1.0.3, isocirc 1.0.7 and long_read_circRNA v2.1 (circNICK-lrs). More information about exact commands and parameters can be found in Supplementary table 4.

### Standardisation of tool output

To ensure comprehensive and standardized evaluation, we converted all tool prediction results to BED12 format, which provides a consistent representation of genomic features including chromosomal location, exon structure, and strand information. This normalization allowed for precise comparative analysis across different circRNA detection tools despite different output formats. More details are given in Supplementary table 5.

### Performance metrics

The bedtools intersect approach utilized three parameters to ensure precise genomic feature comparison. The -f (fraction overlap) parameter controls matching stringency by specifying minimum overlap requirements between predicted and ground truth circRNA annotations. We tested four overlap thresholds: 0.25 (lenient, 25% minimum overlap), 0.5 (moderate, 50% overlap), 0.75 (stringent, 75% overlap), 0.95 (extremely stringent, 95% overlap) to capture a range of detection scenarios.

Our evaluation framework operates at two different levels through the -split and -r parameters. Exon-level evaluation (or block level), enabled by the -split option, treats each BED12 block as a separate feature, testing whether tools can accurately reconstruct complete internal circRNA structure including proper exon boundaries and splice junction patterns. This stringent approach requires precise isoform reconstruction with correct internal splicing. Transcriptome-level evaluation, conducted without -split, assesses overlap at the whole transcript level, regardless of internal exon structure. The -r (reciprocal overlap) parameter ensures symmetric overlap requirements, preventing asymmetric matches where only partial features overlap.

We utilized bedtools version v2.31.1 to compare BED12 format annotations of in silico ground truth against tool outputs. This dual-level approach distinguishes tools that excel at general circRNA boundary detection from those capable of accurate full-length isoform reconstruction.

Performance assessment employed standard metrics across both evaluation levels. True Positives (TP) represent correctly identified circRNAs fitting reciprocal overlap criteria, False Positives (FP) represented computational artifacts or misannotated RNA predictions, and False Negatives (FN) comprised undetected circRNAs from the simulated dataset. We calculated precision, as TP / (TP + FP), that quantifies the proportion of predicted circRNAs that are genuinely correct, effectively measuring the tools’ specificity in avoiding spurious predictions; recall (or sensitivity), calculated as TP / (TP + FN), that assesses the tools’ ability to detect existing circRNAs, revealing their capacity to identify circRNA events within the dataset and the F1 Score, calculated as the harmonic mean of precision and recall, that provides a balanced performance metric and simultaneously considers both false positives and false negatives.

## Results

### Feature extraction from intersected databases and real data for the generation of realistic simulated datasets results

Our circRNA simulation pipeline integrates Nanopore sequencing data from Zhang et al., database annotations, and computational modeling. (Fig. 4).

Among 11,791 circRNAs common to the two databases, canonical GT-AG splice sites dominated the junction boundaries of exonic, exon-intronic, and intronic circRNAs (100%, 99.8%, and 91.2%, respectively), while intergenic circRNAs showed significantly higher usage of non-canonical splice sites (49.5%) (Fig. 3A).

**Figure 3.**
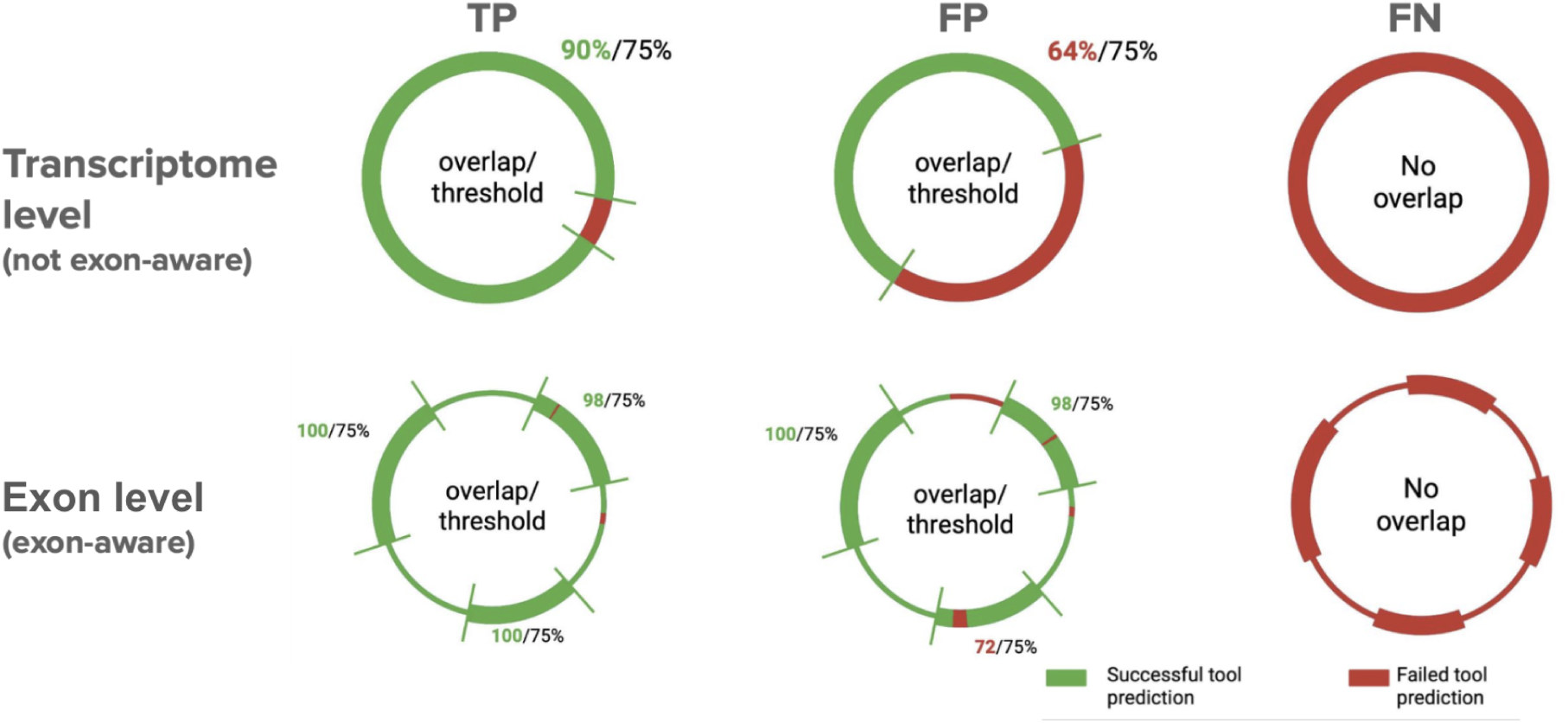
Overlap requirements between predicted and ground truth circRNA annotations on reciprocal overlap threshold 0.75 on transcriptome (not exon aware) and block (exon-aware) levels. TP stands for True Positive, FP for False Positive and FN for False Negative.

**Figure 4.**
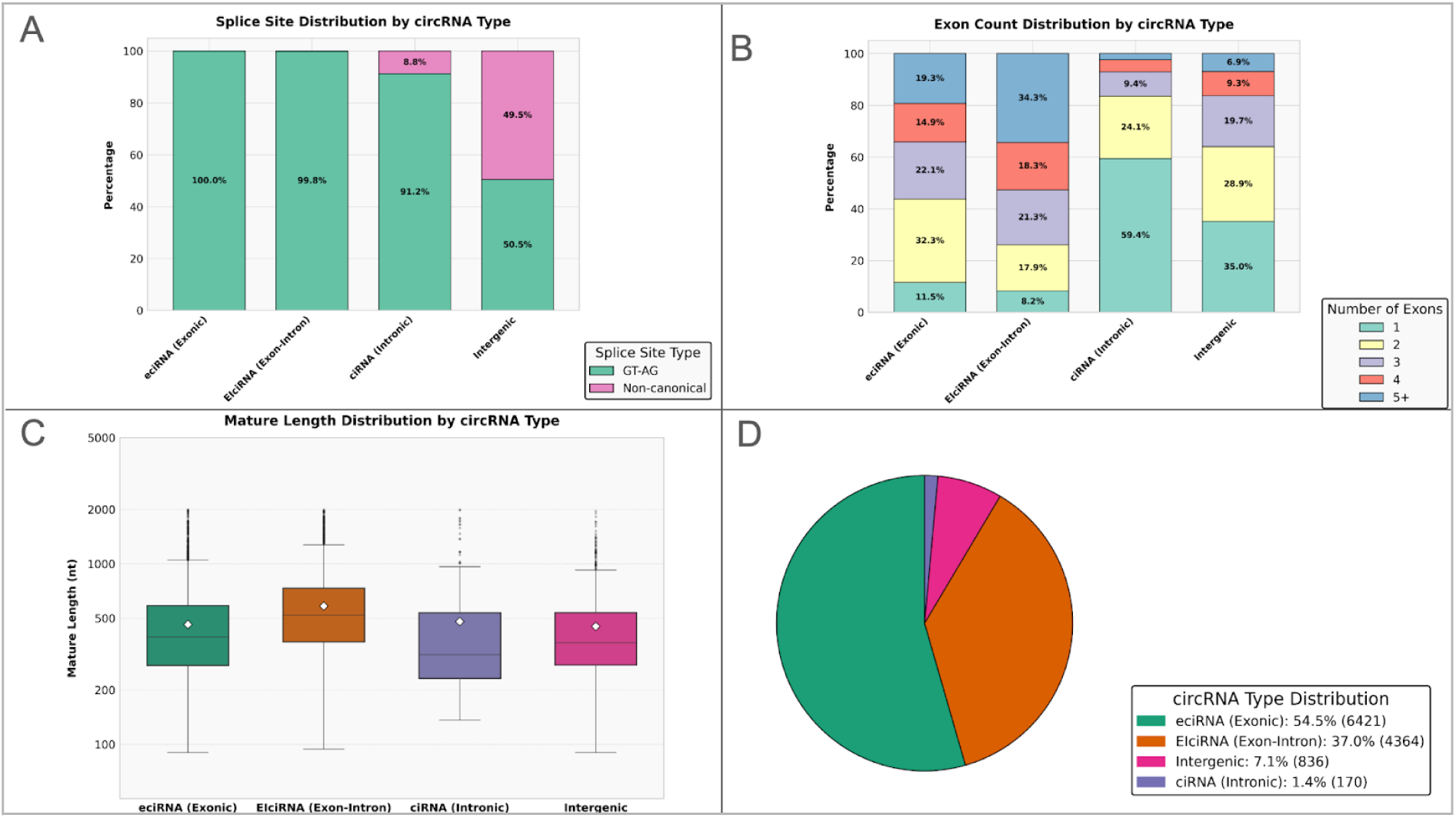
Characterization of Circular RNA molecular features across circRNA types from intersection of mouse circRNAs in CircAtlas and CircBase databases (n=11,791). Panels A-D depict: (A) Splice site composition across circRNA types, (B) Exon count distribution, (C) mature length (exonic concatenation) distribution, and (D) circRNA type distribution.

The exon composition analysis demonstrated substantial variation in complexity across circRNA types (Fig. 3B). Notably, intronic circRNAs (ciRNAs) were predominantly single-exon structures (59.4%), whereas exonic circRNAs (ecircRNAs) showed more diverse exon count distributions with significant proportions containing 2 exons (32.3%) and 3 exons (22.1%). Exon-intron circRNAs (EIciRNAs) exhibited the highest complexity, with 34.3% containing 5 or more exons. Intergenic circRNAs showed an intermediate distribution pattern, with substantial representation across various exon count categories.

Mature length distribution analysis revealed that EIciRNAs possessed the highest median length, followed by ecircRNAs, intergenic, and ciRNAs (Fig. 3C). This pattern reflects the fundamental structural differences between these circRNA subtypes, with EIciRNAs incorporating both exonic and intronic sequences, contributing to their increased overall length.

When examining the overall distribution of circRNA types in the reference databases (Fig. 3D), ecircRNAs constituted the majority (54.5%, n=6421), followed by EIciRNAs (37.0%, n=4364), intergenic circRNAs (7.1%, n=836), and ciRNAs (1.4%, n=170). This distribution highlights the predominance of exon-derived circular RNAs in the current reference datasets and reflects potential biases in detection methodologies favoring exonic circRNA identification.

These different features were incorporated in our simulation framework based on NanoSim to generate circRNA molecules reflecting biological variations. We verified that the simulated data accurately reflected the characteristics of the input parameters. Specifically, the circRNA type splicing patterns, and repeat number variability observed in real Nanopore data and database annotations were preserved in the simulated output.

### Descriptive comparison of circRNA outputs across tools

We performed a comparison of three circRNA detection tools (CIRI-long, isoCirc, and circNick-Irs) run three times on a simulated dataset (n_circRNAs_ = 7503), to evaluate their performance characteristics and detection biases in several modes - with equal proportion of circRNA and linearRNA reads (50%/50% - 200 000 reads each) and realistic proportion expected after circRNA enrichment with only 3% of circRNA reads (Fig. 5) as well as lower (50%/50% - 100 000 reads each) and higher (50%/50% - 300 000 reads each) sequencing depth (Supplementary Fig. 3-5). Specific circRNA simulated, their actual expression, long-read structure and quality varied from run to run. We want to clarify that our descriptions in the results section are based on the 75% reciprocal overlap threshold, which we consider more meaningful for circRNA detection evaluation without sacrificing too much sensitivity. The 75% threshold ensures substantial overlap between predicted and ground truth circRNAs, providing a stringent test of detection accuracy.

**Figure 5.**
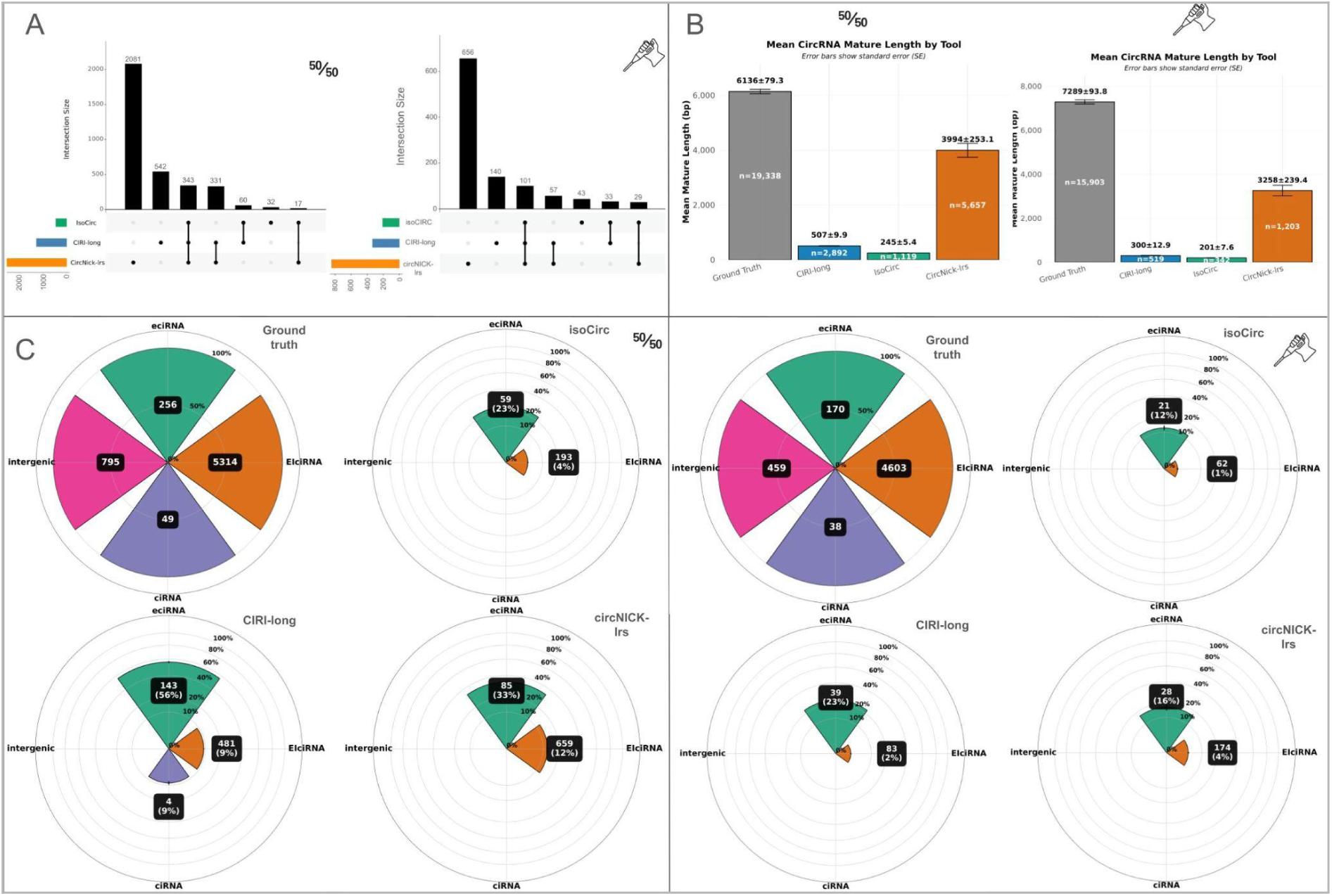
Descriptive comparison of circRNA outputs across tools in 50/50 and realistic modes. Panels A-C depict: (A) intersection analysis showing shared and unique isoform detection across tools. The intersection of circRNA between tools was performed using bedtools intersect with the -split option and an overlap criterion parameter (-f 0.95). The vertical bars represent the number of circRNAs detected in each intersection, and the connected dots below the bars indicate which tool or combination of tools contributes to each intersection, (B) barplots of circRNAs mature length distribution found by each tool, and (C) proportional distribution of circRNA subtypes identified by each tool compared to ground truth.

The intersection analysis revealed significant differences in detection capability among the three tools (Fig. 5A). Notably, circNick-Irs demonstrated the highest unique detection capability, identifying the highest amount of circRNAs not found by other methods - 2081 circRNAs in 50/50 mode and 656 circRNAs in realistic. CIRI-long uniquely detected 542 circRNAs and 140 circRNAs respectively, while isoCirc showed the lowest unique detection with only 32 circRNAs in 50/50 mode and 43 circRNAs in realistic mode. The consensus between all three tools was limited to 434 and 101 circRNAs (10% and 10.83% of total detections). Similar results were obtained in high and low sequencing depth modes, highlighting the complementary nature of these detection approaches. (Supplementary fig. 2)

Analysis of mature circRNA length distributions revealed clear tool-dependent differences in both evaluation modes (Fig. 5B). In the 50/50 mode, CircNick-Irs showed the strongest ability to detect long circRNAs, with a mean length of 4,391 bp, a median of 627 bp, and a very large standard deviation of 22,536 bp, reflecting a highly skewed distribution with an extensive long-length tail and numerous high-length detections. In contrast, CIRI-long primarily detected shorter circRNAs, with a mean length of 504 bp, a median of 297 bp, and a standard deviation of 534 bp, indicating limited representation of longer molecules. IsoCirc exhibited the most restricted length profile, with a mean of 247 bp, a median of 181 bp, and a standard deviation of 177 bp, consistent with its strong bias toward short circRNAs.

A similar pattern was observed in the realistic mode, though with reduced variance across tools. CircNick-Irs continued to outperform other methods in identifying longer circRNAs, with a mean length of 3,258 bp, a median of 933 bp, and a standard deviation of 8,302 bp, maintaining a broad distribution extending into higher length ranges. CIRI-long again showed an intermediate but short-biased profile (mean 300 bp, median 201 bp, standard deviation 293 bp), while IsoCirc remained the most constrained tool (mean 201 bp, median 152 bp, standard deviation 141 bp).

Overall, across both modes, CircNick-Irs consistently demonstrated superior sensitivity to long circRNAs, whereas CIRI-long and IsoCirc were largely limited to short-length detections. The restriction observed for IsoCirc can be attributed to its built-in length cutoff at 4,000 nt in the TRF (Tandem Repeat Finder) section, making it incapable of detecting longer circRNAs without modifying the original code.

These tool-specific length distribution patterns were consistent across both low-depth and high-depth sequencing modes, with sequencing depth primarily affecting detection counts rather than altering the relative mean, median, or dispersion of circRNA lengths for each tool. (Supplementary fig. 4)

When examining the proportional distribution of circRNA types (Fig. 5C), we observed substantial variation in type bias across tools compared to ground truth in both simulation modes. Ground truth composition for the 50/50 mode had a distribution of 82.8% EIciRNAs, 4.0% ecircRNAs, 0.8% ciRNAs, and 12.4% intergenic circRNAs. In the realistic mode, this shifted toward a higher proportion of EIciRNAs (87.3%) and a lower representation of ecircRNAs (3.2%) and intergenic circRNAs (8.7%).

All three tools showed significant detection biases. CircNick-Irs demonstrated the strongest bias toward EIciRNAs, which accounted for 88.6% (50/50) and 86.1% (realistic) of its detections, with limited ecircRNA detection (11.4% and 13.9%, respectively). In contrast, CIRI-long and isoCirc showed a higher detection of ecircRNAs compared to the ground truth proportions. CIRI-long’s detections included 22.8% ecircRNAs in 50/50 mode and 31.9% in realistic mode, while isoCirc showed a similar profile with 23.4% and 25.6% respectively.

Notably, intergenic circRNAs were missed by all three tools in all modes despite representing 12.4% and 8.7% of the ground truth. Similarly, ciRNAs were detected only by CIRI-long in the 50/50 mode (9% of ground truth circRNA), and were entirely missed by all tools in the realistic mode. These findings highlight the importance of tool selection based on the specific circRNA types of interest and suggest that combining multiple tools may provide the most effective circRNA detection across different types.

Across high- and low-depth sequencing conditions, the same classification patterns were preserved. Increased depth raised absolute detection counts but did not meaningfully alter the proportional distribution of circRNA types for any tool. Overall, CircNick-LRS favors EIciRNA detection, CIRI-long and IsoCirc are relatively enriched for ecircRNAs, CIRI-long was the only tool sensitive towards ciRNA and none of the evaluated tools reliably detect intergenic circRNAs. These findings underscore the importance of tool choice when specific circRNA subclasses are of interest and suggest that no single method currently provides comprehensive coverage across all circRNA types. (Supplementary fig. 5)

### Performance evaluation of circRNA detection tools

To evaluate the three circRNA detection tools, we assessed their performance across multiple dimensions including detection accuracy, expression profiling, computational efficiency, and resource requirements (Fig. 6).

**Figure 6.**
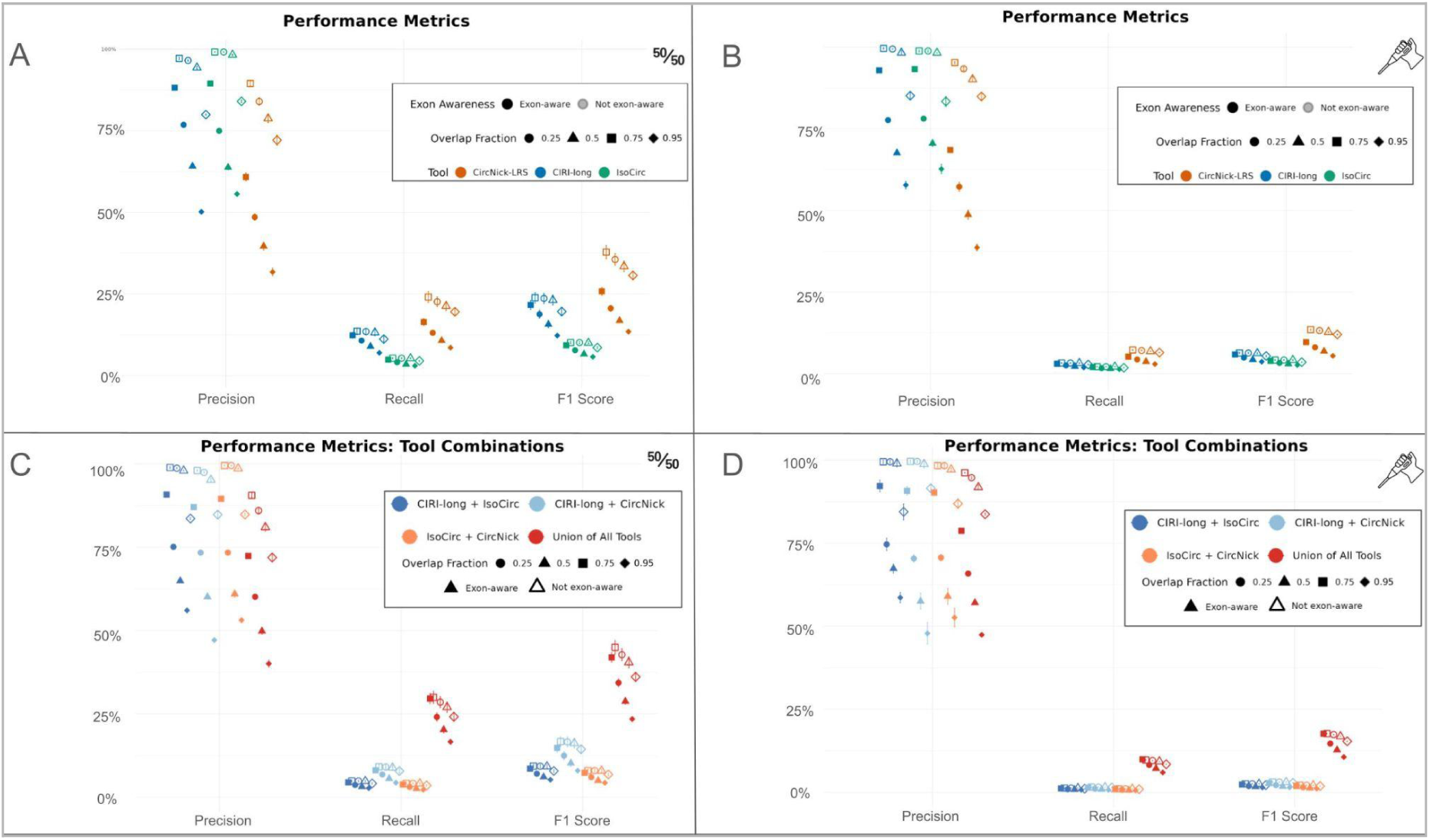
Performance evaluation of circRNA detection tools on simulated datasets in 50/50 and realistic modes. Panels A-D depict: (A-B) quantitative comparison of performance metrics, (C-D) quantitative comparison of performance metrics on combinations and union of tools.

Performance evaluation based on precision, recall, and F1 score revealed tool-specific tradeoffs that were consistent across simulation modes but strongly modulated by evaluation strictness (Fig. 6A–D).

In the 50/50 mode under transcript-level matching (--split off), IsoCirc exhibited the highest precision, with a mean and median of 0.983 and low variability (SD 0.0037), but this came at the cost of very low recall (mean and median 0.055, SD 0.0016), resulting in a modest F1 score (mean 0.105, median 0.104, SD 0.0029). CIRI-long maintained consistently high precision (mean 0.943, median 0.945, SD 0.0067) while achieving higher recall than IsoCirc (mean 0.142, median 0.141, SD 0.0007), yielding an intermediate F1 score (mean and median 0.246, SD 0.0012). CircNick-LRS showed the highest sensitivity, with the greatest recall (mean 0.227, median 0.228, SD 0.0014), albeit with lower precision than the other tools (mean 0.776, median 0.778, SD 0.0042), resulting in the highest overall F1 score in this mode (mean 0.352, median 0.353, SD 0.0021).

In the realistic mode, overall performance declined, primarily driven by reduced recall across all tools, reflecting a more challenging detection environment. IsoCirc retained extremely high precision (mean and median 0.982, SD 0.0010), but recall dropped further (mean and median 0.021, SD 0.0013), producing the lowest F1 score (mean 0.042, median 0.040, SD 0.0025). CIRI-long again showed high precision (mean 0.983, median 0.984, SD 0.0118) with slightly improved recall relative to IsoCirc (mean 0.032, median 0.031, SD 0.0016), yielding an F1 score of mean 0.062 (median 0.061, SD 0.0030). CircNick-LRS remained the most sensitive tool, achieving the highest recall in this mode (mean 0.069, median 0.068, SD 0.0013), though precision decreased relative to 50/50 mode (mean 0.901, median 0.907, SD 0.0132), resulting in the best F1 score among tools (mean 0.127, median 0.127, SD 0.0021). Across both modes, the relative ranking of tools was preserved: IsoCirc maximized precision, CircNick-LRS maximized recall, and CIRI-long occupied an intermediate position.

Applying exon-level matching (--split on) substantially changed performance results in both modes, highlighting the difficulty of reconstructing exact internal exon structures. In the 50/50 mode at 0.75 overlap, mean precision dropped sharply for all tools: from 0.776 to 0.385 for CircNick-LRS, from 0.943 to 0.640 for CIRI-long, and from 0.983 to 0.633 for IsoCirc, with corresponding reductions in F1 scores. A similar pattern was observed in the realistic mode, where mean precision decreased from 0.901 to 0.487 for CircNick-LRS, from 0.983 to 0.676 for CIRI-long, and from 0.982 to 0.705 for IsoCirc. These declines reflect inner exon-boundary mismatches rather than failures in circRNA detection, indicating that many transcript-level matches do not reproduce the correct internal splicing architecture.

Combining predictions from multiple tools further emphasized the distinction between transcript-level detection and isoform-level correctness. In the 50/50 mode without exon-level enforcement, the union of all three tools produced the highest recall (mean and median 0.366, SD 0.002), substantially substantially outperforming any individual tool or pairwise combination, but with reduced precision (mean 0.845, median 0.846, SD 0.004). Enabling --split caused a decline in performance, with mean precision dropping to 0.489 and recall to 0.212, leading to a substantial reduction in F1. In the realistic mode, the union strategy again maximized sensitivity under transcript-level matching (mean recall 0.121, median 0.120, SD 0.001) while maintaining high precision (mean and median 0.941, SD 0.002). However, enforcing exon-level matching reduced mean precision to 0.580 and recall to 0.062, confirming that structural inconsistencies accumulate rather than resolve when combining tools.

Across low- and high-depth sequencing conditions (Supplementary fig. 6), the qualitative trends remained unchanged. Increased depth primarily improved recall, while precision was comparatively stable, and the relative ranking of tools and combinations was preserved.

Collectively, these results demonstrate that while tool choice and combination strategies strongly influence detection sensitivity, accurate reconstruction of internal circRNA isoform structure remains a major computational challenge, particularly under realistic and exon-level evaluation constraints.

### Expression analysis

Expression analysis revealed clear differences in how tools recover circRNA abundance and how detection sensitivity varies across expression levels (Fig. 7). In the 50/50 mode, IsoCirc demonstrated the strongest agreement with ground-truth expression among detected circRNAs, with a Pearson correlation of r = 0.779 (R² = 0.606) and a Spearman correlation of 0.727, indicating reliable recovery of both absolute expression trends and relative ranking. CIRI-long showed slightly lower but still strong agreement (Pearson r = 0.729, R² = 0.532; Spearman 0.761), whereas CircNick-LRS showed weaker quantitative agreement (Pearson r = 0.483, R² = 0.233; Spearman 0.367), despite detecting a larger number of circRNAs. This indicates that CircNick-LRS favors sensitivity over quantitative accuracy, while IsoCirc and CIRI-long better preserve expression proportionality.

**Figure 7.**
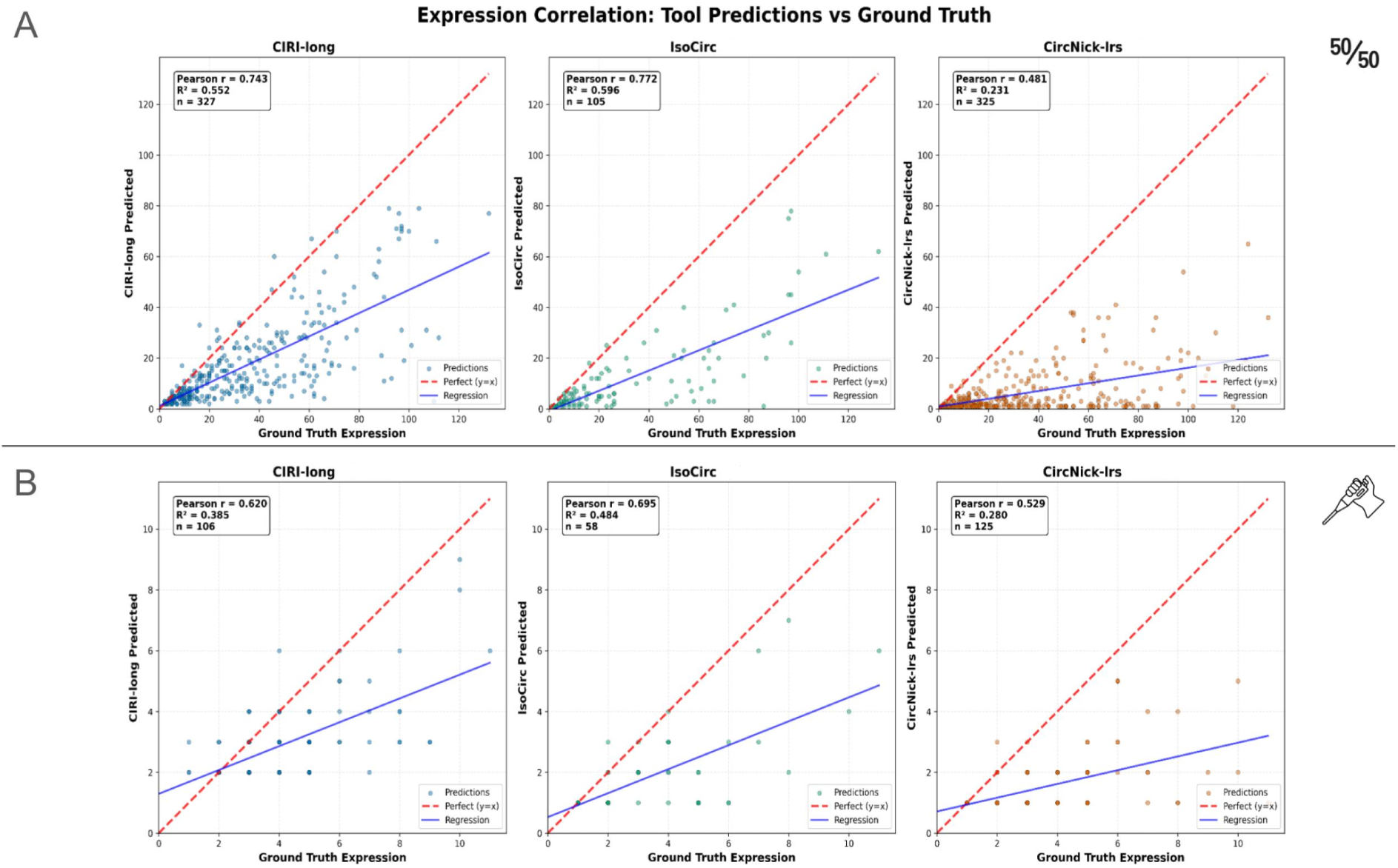
Correlation between predicted and ground-truth circRNA expression. Scatter plots show predicted circRNA expression versus ground-truth expression. The dashed red line shows perfect agreement (y = x), and the solid blue line shows the linear regression fit. Pearson correlation coefficients (r), coefficients of determination (R²), and the number of circRNAs (n) are reported in each panel. Panel (A) represents 50/50 mode, while panel (B) - realistic mode.

Stratifying circRNAs by expression level by tertiles revealed distinct sensitivity profiles for each tool in the 50/50 mode. CircNick-LRS showed the highest sensitivity overall and was particularly effective at detecting medium-expression circRNAs, with sensitivities of 0.133 (low), 0.299 (medium), and 0.240 (high). CIRI-long showed intermediate sensitivity, again peaking in the medium-expression group (0.070 low, 0.199 medium, 0.153 high), while IsoCirc showed the lowest sensitivity across all expression levels (0.036 low, 0.072 medium, 0.058 high), reflecting a conservative detection strategy. Across all tools, medium-expression circRNAs were consistently the most detectable. (Supplementary fig. 7)

In the realistic mode, expression concordance declined for all tools, reflecting increased transcriptome complexity. IsoCirc continued to show the strongest quantitative agreement (Pearson r = 0.695, R² = 0.484; Spearman 0.629), followed by CIRI-long (Pearson r = 0.620, R² = 0.385; Spearman 0.540), while CircNick-LRS remained the weakest in expression accuracy (Pearson r = 0.529, R² = 0.280; Spearman 0.539). Sensitivity profiles in this mode showed a collapse for high-expression circRNAs, which were rarely detected by any tool. CircNick-LRS retained the highest sensitivity for low-and medium-expression circRNAs (0.087 low, 0.120 medium), whereas CIRI-long (0.020 low, 0.081 medium) and IsoCirc (0.028 low, 0.036 medium) showed more limited detection. Across all tools, medium-expression circRNAs again represented the most detectable class, highlighting a consistent bias toward intermediate abundance levels. (Supplementary fig. 7)

Across high- and low-depth sequencing conditions (Supplementary fig. 7-8), these expression-dependent sensitivity patterns remained stable. Increased sequencing depth improved sensitivity across all expression strata, particularly for medium-expression circRNAs, but did not alter the relative behavior of the tools. Overall, CircNick-LRS is best suited for detecting a broad range of circRNAs, particularly those with moderate expression, CIRI-long provides balanced performance with moderate sensitivity and strong expression concordance, and IsoCirc prioritizes accurate detection and quantification at the expense of sensitivity, especially for low-abundance circRNAs.

### Computational performance

Resource utilization varied dramatically between tools (Fig. 8), reflecting architectural differences observed under identical testing conditions with providing 8 threads and identical input datasets. CIRI-long exhibited exceptionally high memory consumption (307.07 ± 27.93 GB), representing a substantial computational burden. This likely comes from its approach to simultaneous processing of multiple consensus sequences per thread, maintenance of large genome index structures for detecting repetitive patterns, and rolling circle detection algorithms that require extensive k-mer matching. Notably, this memory consumption pattern suggests that providing too many threads to CIRI-long could lead to OOM (Out of Memory) errors during a run; thus, it is recommended to limit available cores for this tool.

**Figure 8.**
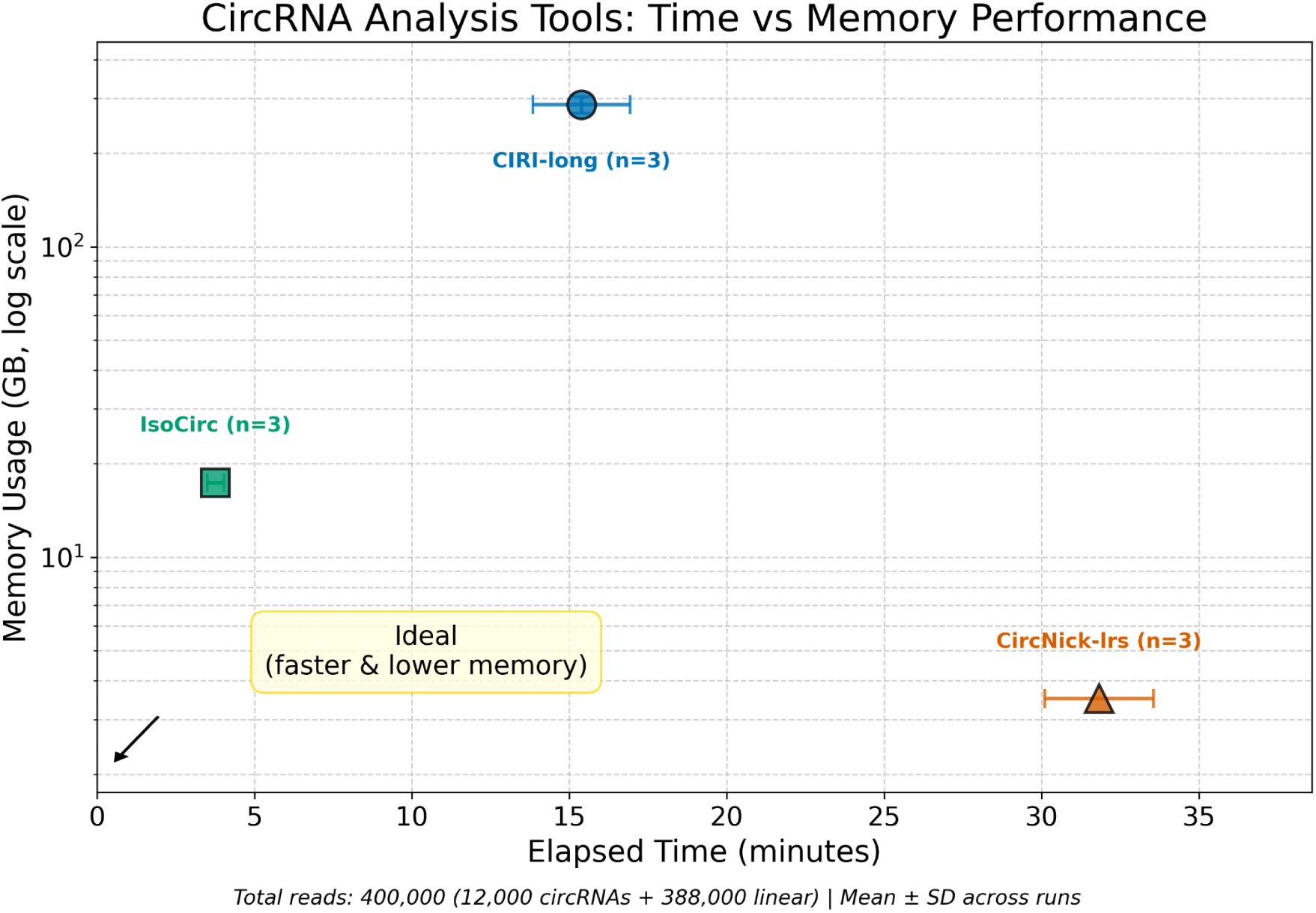
Computational performance and resource usage. Dot plot showing peak memory usage requirements for each computational approach and displaying processing time per read when analyzing a standardized dataset of 400,000 reads on 8 provided threads.

In contrast, isoCirc (17.37 ± 0.06 GB) demonstrated a memory-efficient design with remarkably low variability, while circNick-LRS (3.47 ± 0.06 GB) showed the most efficient memory utilization overall. Importantly, circNick-LRS’s low memory footprint is connected to its single-threaded architecture, which prevents the memory multiplication occurring in multi-threaded processing. This makes both tools highly accessible for researchers with limited computational resources, albeit at the cost of processing speed in circNICK-lrs case.

Processing speed analysis revealed significant performance differences that reflect different computational architectures. IsoCirc emerged as the clear performance leader, achieving a throughput of 6,252,897 ± 637,278 reads per hour - processing our dataset of 400 000 reads in just 3.87 minutes. This demonstrates superior computational efficiency through optimized multi-threading. CIRI-long demonstrated intermediate efficiency, processing 1,507,034 ± 139,648 reads per hour (approximately 16.02 minutes elapsed time for our dataset). Meanwhile, circNick-LRS required the most processing time, with a throughput of 726,393 ± 40,565 reads per hour (33.11 minutes elapsed), making it the slowest of the three tools.

## Discussion

Our benchmarking study reveals the current state of circRNA detection tools for long-read Nanopore sequencing, highlighting both the promising capabilities and significant challenges in accurately identifying circular RNAs. The analysis highlights the complementary strengths and limitations of current computational approaches, providing insights for researchers.

Each tool demonstrated distinct performance characteristics that require consideration.

IsoCirc emerged as the computational efficiency leader, making it an attractive option for researchers requiring rapid analysis, with memory-efficient design and remarkably low variability. IsoCirc excelled at precision and expression quantification accuracy, demonstrating the strongest agreement with ground-truth expression among detected circRNAs, making it the best choice for studies prioritizing precision and more accurate quantification of circRNA abundance. However, this precision and computational efficiency came at the cost of very low recall, resulting in the lowest overall sensitivity among the three tools, with particular weakness in detecting longer circRNAs. Its built-in length cutoff at 4,000 nucleotides in the TRF module restricts detection of longer circRNAs, and it exhibited a notable bias toward detecting ecircRNAs compared to ground truth proportions.

CIRI-long offered a more balanced approach, occupying an intermediate position across most performance metrics, with consistently high precision and higher recall than IsoCirc. Like other tools, CIRI-long completely missed intergenic circRNAs, but it was uniquely capable of detecting ciRNAs, being the only tool to identify three circRNA types. It also showed strong expression concordance. However, CIRI-long imposed a substantial computational burden with exceptional memory consumption - a significant constraint for many research environments - and we would recommend limiting available cores for this tool to avoid out-of-memory errors. The tool also showed limitations in detecting longer circRNAs, and exhibited a relatively high bias toward ecircRNA detection compared to ground truth proportions.

CircNick-LRS distinguished itself by demonstrating the highest overall sensitivity, achieving the highest recall among all methods and the best F1 scores, with the highest number of uniquely identified circRNAs. The tool excelled particularly at detecting longer circRNAs, maintaining a broad distribution extending into higher length ranges, and demonstrated the highest sensitivity across all expression levels. However, CircNick-LRS showed the lowest precision among the three tools and the weakest quantitative agreement with ground-truth expression, and struggled most severely with predicting correct exon structure, showing the sharpest decline in precision when exon-level matching was enforced. Notably, it also exhibited the strongest bias toward EIciRNA detection. And its single-threaded architecture, while memory-efficient, results in the slowest processing speed. The tool is also restricted to built-in mouse and human reference genomes (mm10 and hg19) with predefined annotation files, preventing its use with custom or alternative genome annotations.

The extremely low intersection between tools strongly suggests that relying on a single detection tool is suboptimal - each tool acted as a unique lens, capturing distinct aspects of the circRNA landscape that others missed. All tools also exhibited consistent bias toward detecting medium-expression circRNAs, with lower sensitivity for both low-and high-expression circRNAs, further complicating comprehensive circRNA characterization.The choice of tool or combination should be guided by specific research priorities and practical constraints. For maximum sensitivity, the union of all three tools provides the highest likelihood of detecting true circRNAs, making it valuable for exploratory studies, though at the cost of reduced precision and increased false positives. For accurate expression quantification, IsoCirc or CIRI-long are the preferred choices, with IsoCirc also offering the lowest computational footprint when resources are limited. For balanced detection with reasonable precision and sensitivity, particularly when longer circRNAs are of interest, CircNick-LRS offers the best single-tool compromise in terms of F1 scores, though its single-threaded architecture and longer processing time should be considered for large datasets. When high precision is needed and false positives must be minimized, the combination of CIRI-long and IsoCirc maintains the highest precision values while improving recall over IsoCirc alone. Regardless of tool choice, it is strongly recommended to verify the internal structure of detected circRNAs of interest using orthogonal approaches.

Across these three pipelines, several methodological choices may introduce systematic biases in circRNA detection. Differences in mapping algorithms (minimap2, bwa mem, pblat), filtering criteria (canonical splice signals, mapping quality thresholds, genomic constraints), and validation against existing circRNA databases may introduce systematic biases that preferentially retain well-annotated exon-derived circRNAs while reducing detection of intronic, non-canonical, or intergenic events. This was directly reflected in our results: all three tools completely missed intergenic circRNAs, which represented 12.4% of the ground truth dataset in 50/50 mode and 8.7% in realistic mode, and ciRNAs were detected only minimally by CIRI-long and entirely missed by all tools in the realistic mode. These findings highlight that no single method currently provides comprehensive coverage across all circRNA types, and tool combinations or development of new methodologies are needed to address these blind spots. We hypothesize that intergenic circRNAs escaped detection because tool pipelines were primarily developed and optimized using annotations from known genes, reducing their sensitivity to circRNAs originating from intergenic regions that lack well-defined splice site motifs (Supplementary Figure 1) or reference annotations. In addition, intergenic circRNAs may display distinct sequence characteristics, such as differences in GC content, weaker or non-canonical splice signals, or atypical length distributions. Although our simulation of intergenic circRNAs was guided by parameters extracted from existing databases, it likely does not fully capture their biological complexity, and more advanced approaches such as deep learning-based models may be required to better approximate their properties. Beyond circRNA type-specific detection gaps, sensitivity remained consistently low across all tools when exon-level matching was enforced, with precision dropping sharply compared to transcript-level evaluation. This highlights the challenge of accurately reconstructing exact internal exon structures, with many transcript-level matches failing to reproduce the correct internal splicing architecture. These findings suggest that while tools are reasonably effective at identifying circRNA loci, accurate isoform-level reconstruction remains a major computational challenge requiring further methodological development.

Our results show mixed consistency with the original publications, reflecting differences in evaluation approaches. CIRI-long’s original paper (Zhang et al., 2021) reported higher performance metrics (F1 score of 0.92 for read-level analysis) in simulation studies, but these focused on relatively simple ecircRNA structures generated from inner exons without modeling other circRNA types or complex features like exon skipping or intron retention, and analysis was conducted on read-level, not on exon/transcriptome level approach we employed here. Our more complex simulation framework, incorporating four circRNA types with alternative splicing patterns, may explain the performance differences from their original validation. The isoCirc publication (Xin et al., 2021) used real datasets for validation, demonstrating high accuracy for full-length circRNA reconstruction across 12 human tissues and HEK293 cells, with emphasis on detecting alternative splicing events within circRNAs that aligns with our observation of isoCirc’s precision capabilities. The circNick-LRS study (Rahimi et al., 2021) focused on characterizing circRNA diversity in human and mouse brain samples, with validation primarily through RT-PCR of selected candidates rather than systematic benchmarking metrics. Importantly, none of the prior studies used the similar simulation framework or standardized evaluation metrics across all three tools, making our simulation environment with known ground truth suitable for revealing performance characteristics and tool overlaps that were not systematically evaluated in the original publications.

Limitations of the current study include the exclusion of the circFL-seq tool from the benchmarking analysis. This tool was initially considered for inclusion but was ultimately omitted since we were unable to obtain a functional installation despite following the authors’ instructions. The dataset for this study was generated using CIRI-long dataset parameters, which may introduce some methodological bias. In future versions of this study, we plan to expand our approach by using FASTQ files from different wet-lab protocols as a base for in-silico dataset generation, as well as incorporating various biological sources (mouse brain tissue and HEK293 cell cultures) for a more comprehensive performance evaluation across diverse experimental conditions. This expansion will provide a more robust assessment of tool performance in varied biological contexts and sequencing approaches, further enhancing the utility of our benchmarking framework for the circRNA research community.

Notably, all three circRNA detection tools lacked container environments, posing installation challenges. CIRI-long and isoCirc proved particularly difficult to deploy, requiring multiple reinstallations following cluster updates. To resolve this issue we built containers for both tools, which are provided alongside this manuscript for community use (see Data, scripts, code, and supplementary information availability). We recommend that developers of circRNA detection tools prioritize containerization, as these tools are primarily used by researchers without extensive bioinformatics expertise to analyze their wet-lab data, and installation barriers can significantly limit tool accessibility and adoption.

We also acknowledge that potential biases may arise from our use of existing circRNA databases to guide feature extraction, as many annotated circRNAs were originally identified using short-read sequencing approaches and lack comprehensive experimental validation. Although our benchmark was conducted on mouse-based simulated datasets, extending the framework to human transcriptomes is an important future direction. Differences in genome complexity, annotation depth, and circRNA diversity may influence tool performance, and a dedicated human-specific evaluation would provide complementary insights. Moreover our current pipeline does not systematically generate isoform families, though it does create different circRNAs from the same gene with varying internal structures (but not the same BSJ). As a future perspective we are implementing an isoform mode which would allow us to model isoform families sharing BSJ coordinates but differing in internal structure and alternative splicing events within the same circRNAs boundaries.

In conclusion, this study provides an evaluation of circRNA detection tools, emphasizing the need for continued methodological refinement and strategic tool selection. Researchers must carefully consider the specific requirements of their studies - computational resources, circRNA length, expression levels, circRNA types of interest, and the balance between precision and sensitivity - when selecting circRNA detection tools. By integrating wet-lab data, database annotations, and computational modeling, our framework captures circRNA biogenesis complexity and provides a valuable resource for studying circular RNA function and regulation. Future iterations of this benchmarking study will continue efforts to incorporate additional circRNA detection tools, potentially developing a more standardized approach to tool integration, documentation, and user accessibility. Our scripts are freely available at https://gitlab.com/bioinfog/circall/nano-circ.

## Acknowledgements

Preprint version of this article has uploaded to bioRxiv (https://doi.org/10.1101/2025.04.17.649290) and peer-reviewed and recommended by Peer Community In PCI Genomics (Marchet, C. (2026) Evaluating circular RNA detection in the long-read era. Peer Community in Genomics, 100439. https://doi.org/10.24072/pci.genomics.100439).

The authors would like to thank the Gene Expression and Oncogenesis team, the Dog Genetics team and Biology in Silico 2.0 group (IGDR CNRS UMR6290) for helpful discussions. We also acknowledge the GenOuest bioinformatics core facility (https://www.genouest.org/) for providing the computing infrastructure.

## Data, scripts, code, and supplementary information availability

Data are available online: https://doi.org/10.1038/s41587-021-00842-6 (Zhang et al., 2021, https://doi.org/10.17605/OSF.IO/76FGW (Rusakovich et al., 2026);

Scripts and code are available online: https://doi.org/10.5281/zenodo.15229596 (Rusakovich et al, 2025);

Containers for circRNA analysis tools are available online: https://doi.org/10.5281/zenodo.18669388 (Rusakovich et al., 2026).

### Supplementary materials

**Supplementary Figure 1.**
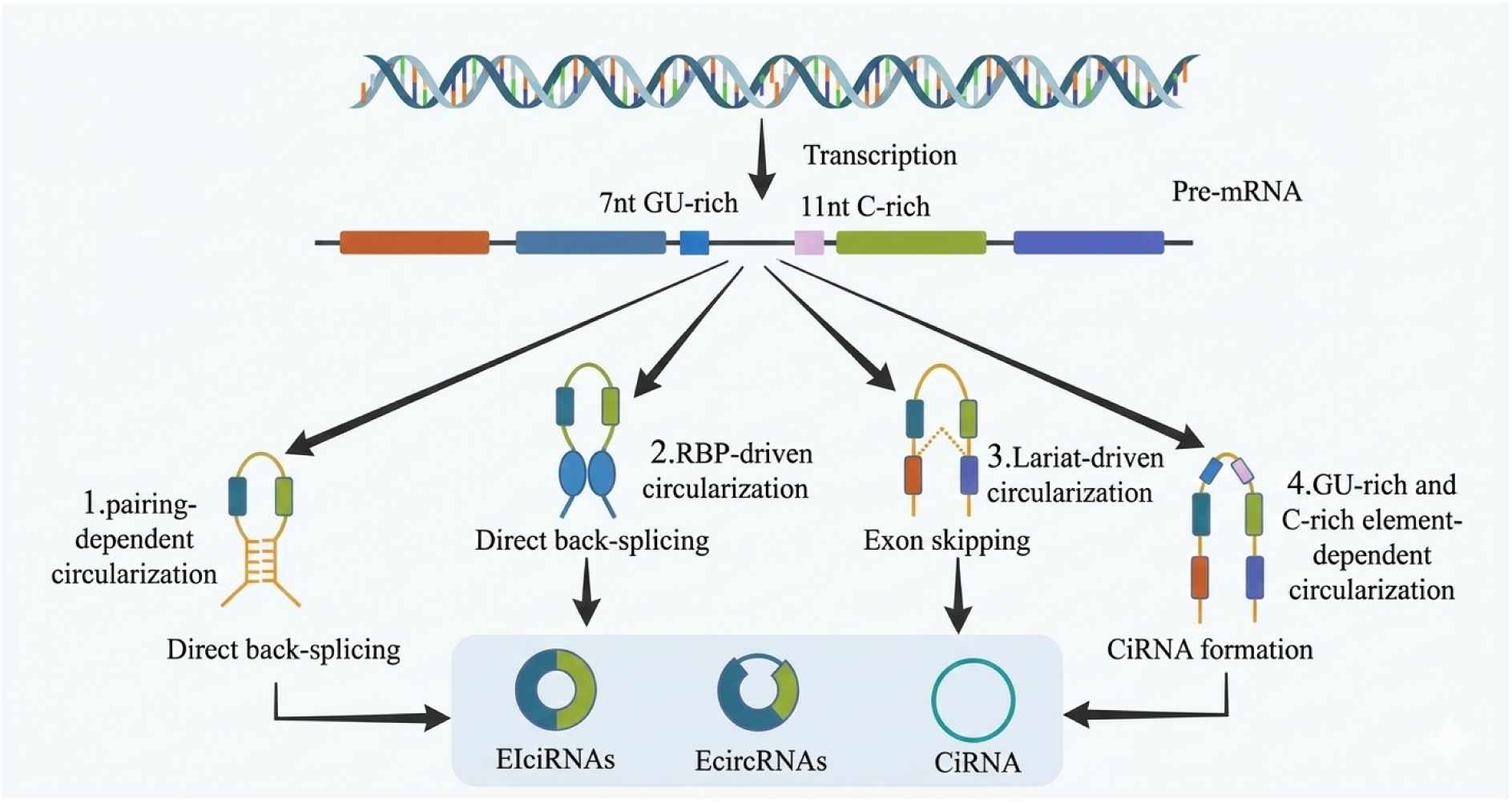
Types of circular RNAs and their biogenesis. Schematic overview of circular RNA (circRNA) types generated from pre-mRNA through different back-splicing mechanisms. circRNAs are classified into exonic circRNAs (eciRNAs), composed exclusively of exons; exon-intron circRNAs (EIciRNAs), which retain intronic sequences between circularized exons; and circular intronic RNAs (ciRNAs), derived solely from intronic regions. Figure adapted from Ma Y, Zheng L, Gao Y, Zhang W, Zhang Q, Xu Y. A Comprehensive Overview of circRNAs: Emerging Biomarkers and Potential Therapeutics in Gynecological Cancers. Front Cell Dev Biol. 2021.

**Supplementary Figure 2.**
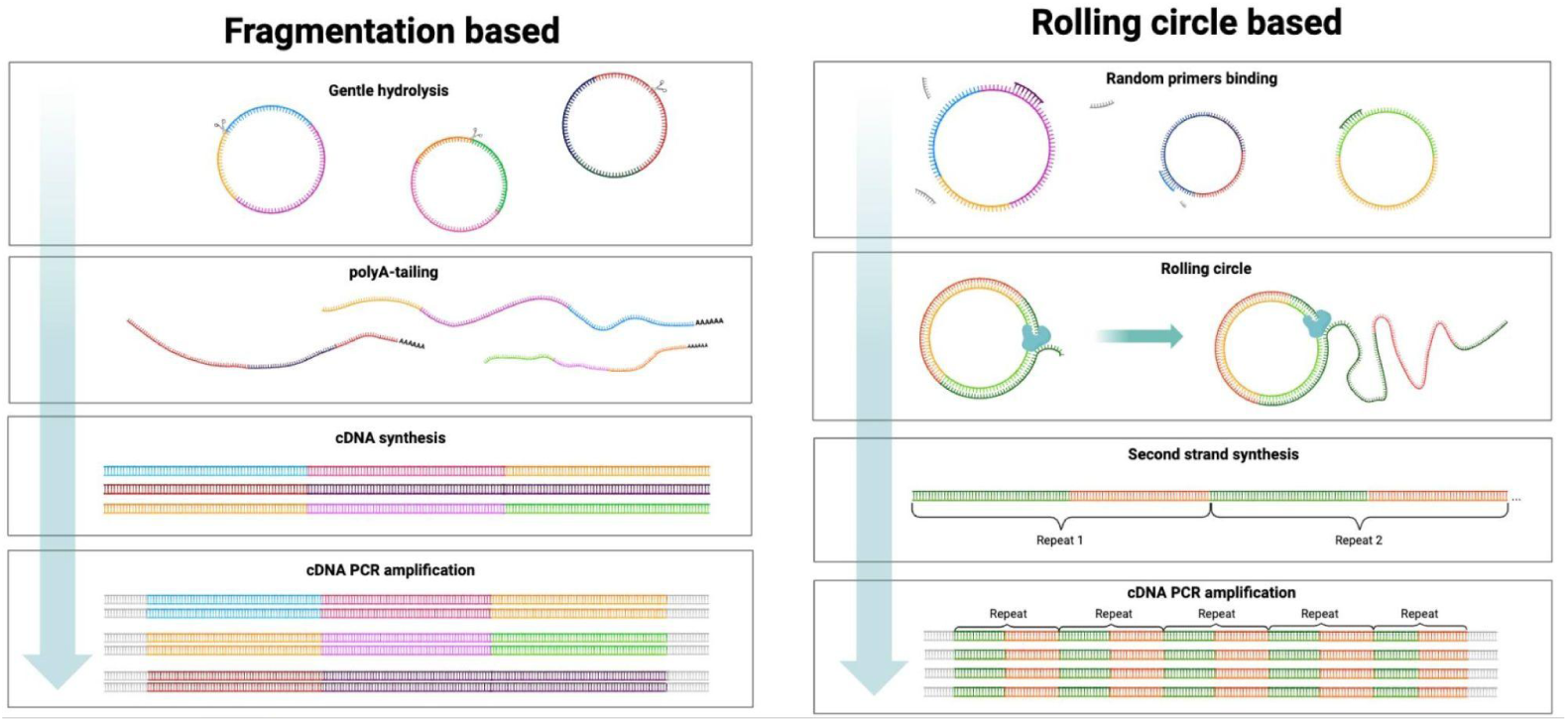
Expected sequence based on wet-lab approach. (A) Nicking approach utilised by circNICK-lrs tool provides one circRNA - one read pattern. (B) Rolling circle approach utilised by CIRI-long and isoCIRC have several repeats of circRNA per read creating during rolling circle reverse transcription or rolling circle amplification step.

**Supplementary Figure 3.**
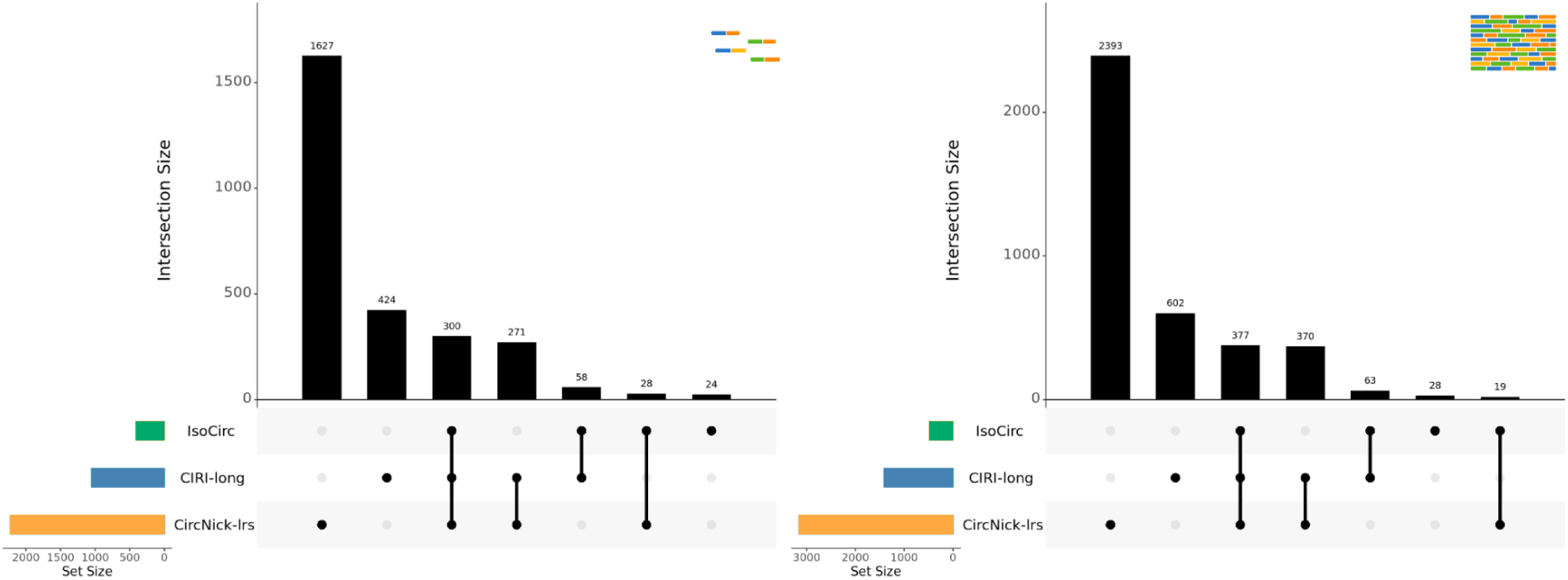
intersection analysis showing shared and unique isoform detection across tools in lower depth (100 000 reads per RNA type) vs higher depth (300 000 reads per RNA type) modes.

**Supplementary Figure 4.**
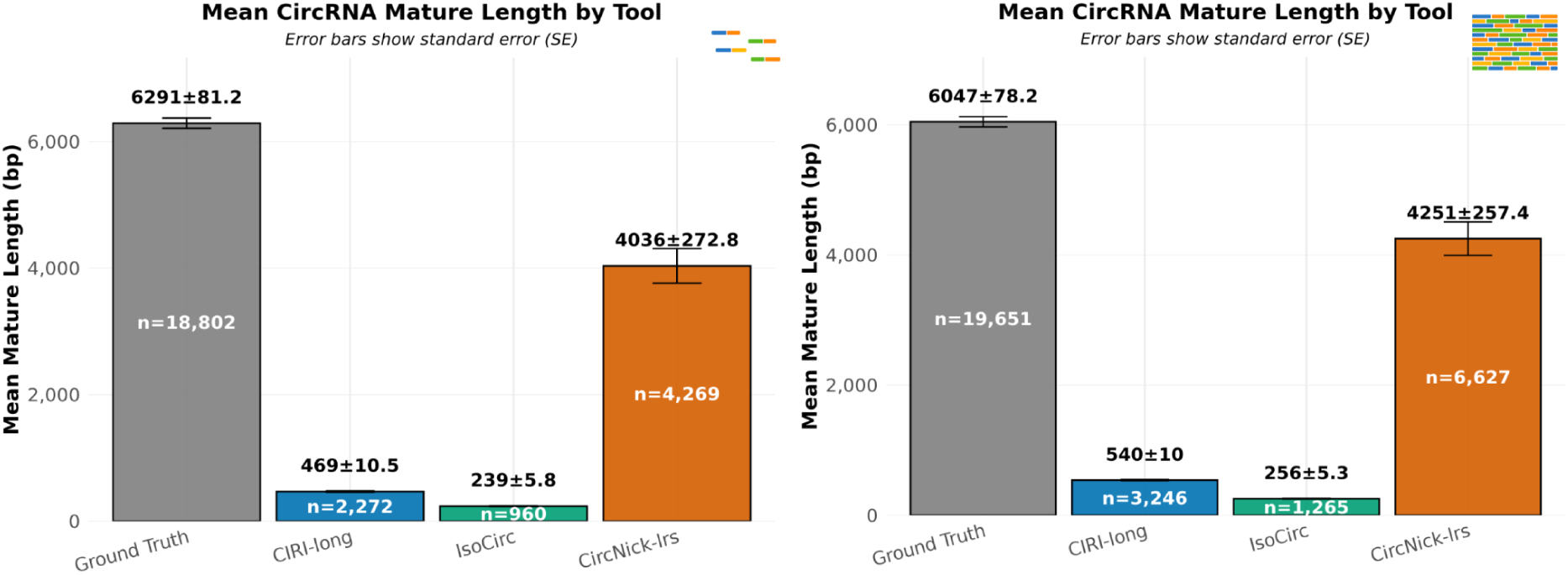
circRNAs mature length distribution found by each tool in lower depth (100 000 reads per RNA type) vs higher depth (300 000 reads per RNA type) modes.

**Supplementary Figure 5.**
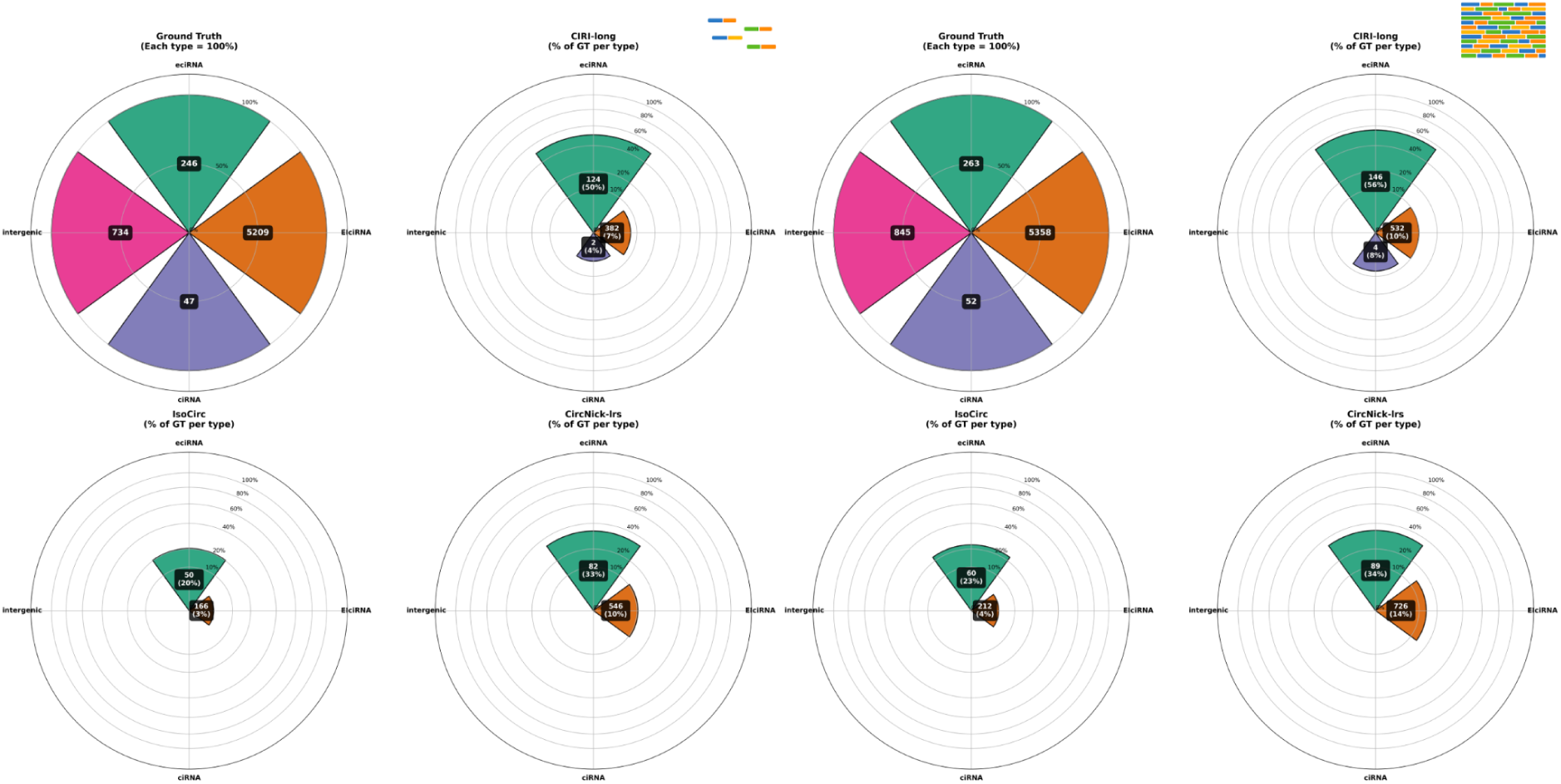
proportional distribution of circRNA subtypes identified by each tool compared to ground truth in lower depth (100 000 reads per RNA type) vs higher depth (300 000 reads per RNA type) modes.

**Supplementary Figure 6.**
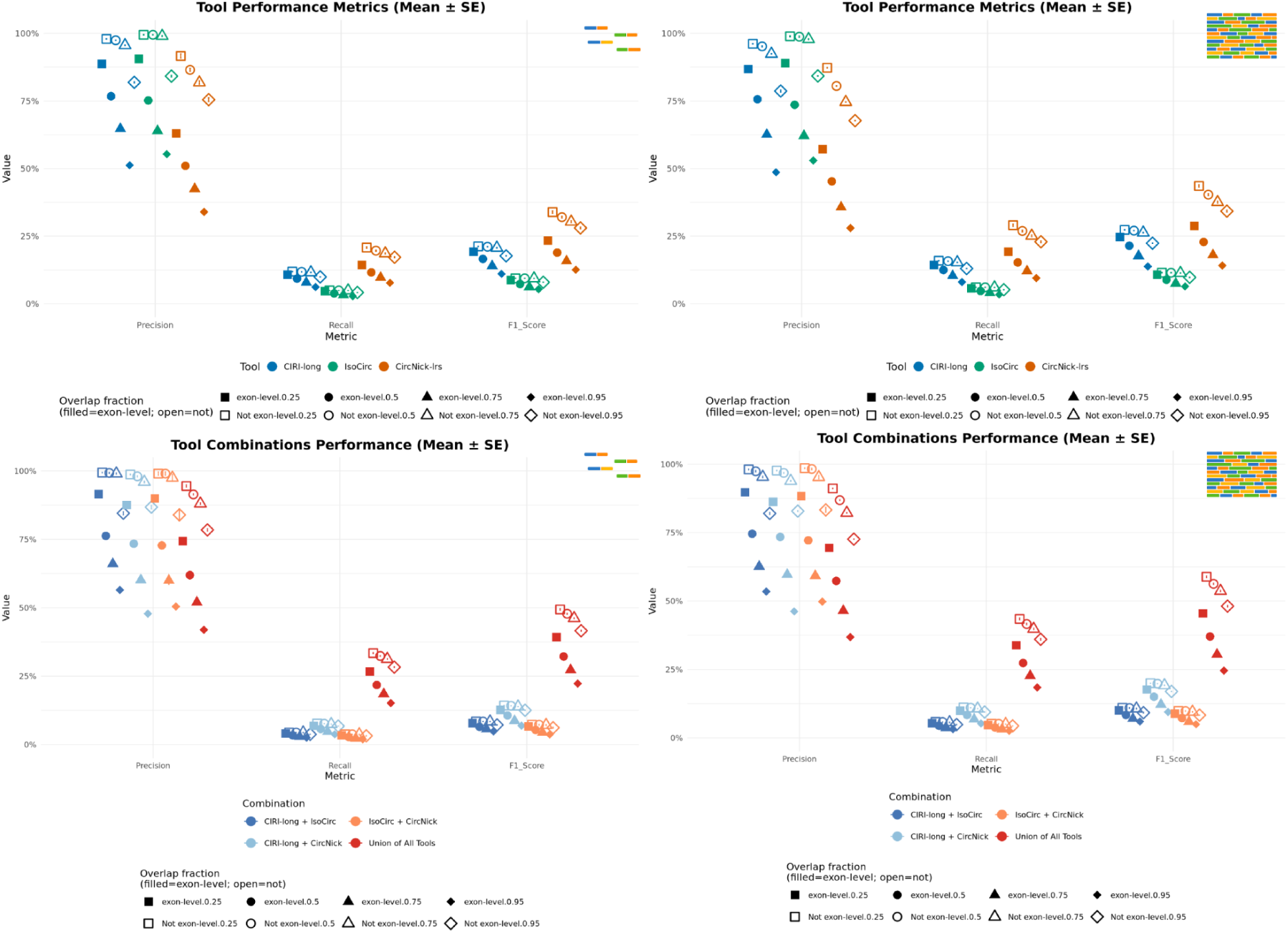
quantitative comparison of performance metrics for tools, combinations and union of tools in lower depth (100 000 reads per RNA type) vs higher depth (300 000 reads per RNA type) modes.

**Supplementary Figure 7.**
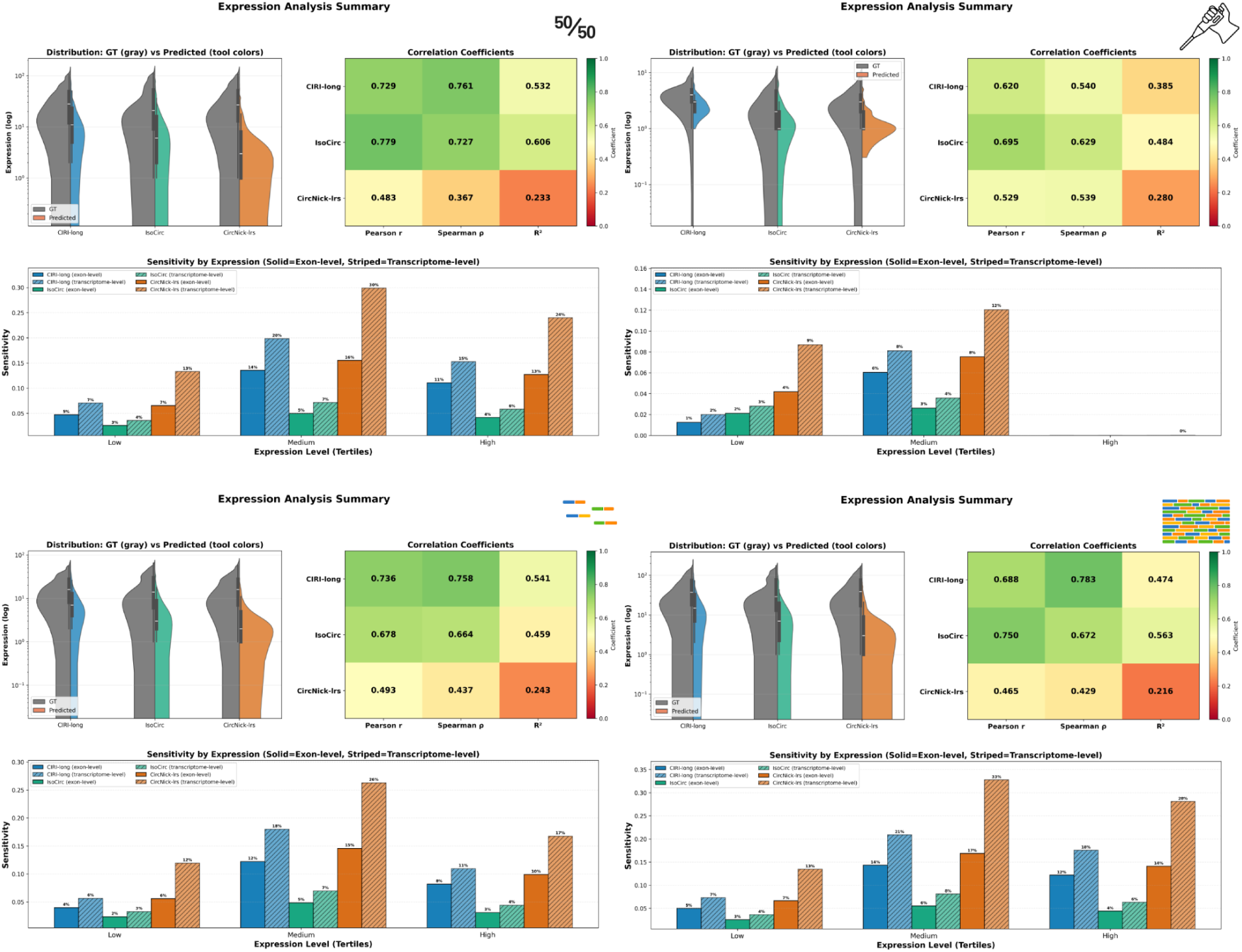
expression prediction performance under different modes. Comparison of 50/50, realistic, lower depth (100 000 reads per RNA type), and higher depth (300 000 reads per RNA type) modes using distributional agreement, correlation coefficients, and expression-stratified sensitivity.

**Supplementary Figure 8.**
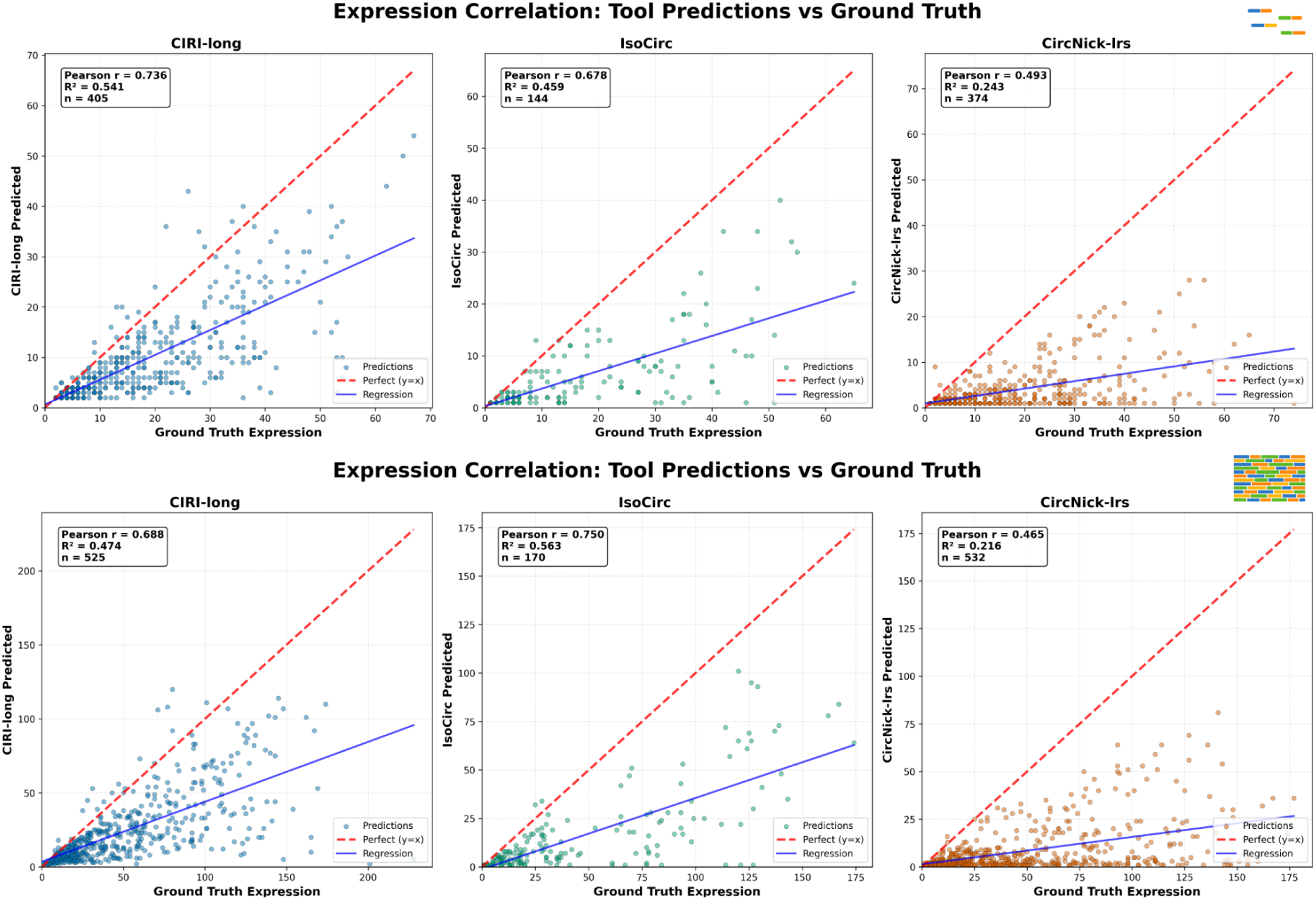
Correlation between predicted and ground-truth circRNA expression. Scatter plots show predicted circRNA expression versus ground-truth expression. The dashed red line shows perfect agreement (y = x), and the solid blue line shows the linear regression fit. Pearson correlation coefficients (r), coefficients of determination (R²), and the number of circRNAs (n) are reported in each panel. The top panel represents low-depth mode, while the bottom panel - high-depth mode.

**Supplementary Table 1.** Comparison of key features and computational components across three long-read circular RNA sequencing methods.

| Feature | CIRI-long | IsoCirc | circNick-LRS |
| --- | --- | --- | --- |
| Input | RCRT-based ONT reads | RCA-based ONT reads | Linearised circRNA ONT reads |
| Mapping tool | mappy (v2.17) | minimap2 (v2.17) | pblat (v35) |
| Circular pattern detection | k-mer matching | Tandem Repeat Finder (v 4.0.9) | - |
| **BSJ detection** | SPOA,<br>mappy (v2.17), bwapy<br>(v0.1.4) | minimap2 (v2.17) | pblat (v35),<br>bedtools (v2.29.2) |
| **Reference**<br>**databases** | circAtlas | circBase, MiOncoCirc | circBase,<br>circAtlas,<br>CIRCpedia |

**Supplementary Table 2.** Features that are used in circRNA simulation.

| **Parameter** | **Description / Purpose** | **Source** |
| --- | --- | --- |
| **Number of circRNAs** | Total amount of circRNAs to generate | User defined |
| **Proportion of circRNA type** | eciRNA, ciRNA, ElciRNA, intergenic | User defined |
| **Mean copy number (13.50) of rolling circles** | Used as the basis for copy number distribution. | Wet-lab derived |
| **CircRNA length distribution** | Used for length distributions by type. | Database derived |
| **Gene and transcript IDs** | Retrieved from Ensembl annotations. | Database derived |
| **CircRNA length ranges** | Constrains intergenic circRNA sizes. | Database derived |
| **Exon block structure** | Generated based on exon count distributions by type. | Database derived |
| **Exon count distributions by type** | Includes proportions for 1, 2, 3, 4, and 5+ exon structures for different circRNA types. | Database derived |
| **Splice site type by circRNA type** | GT-AG, GC-AG, AT-AC, and non-canonical splice types. | Database derived |
| **Copy number distribution ranges** | Defines realistic ranges for simulation. | Randomised |
| **Chromosome, start, end coordinates** | Generated for each simulated circRNA. | Randomised (by finding suitable splice site) |
| **Strand orientation** |  | Randomised |
| **TPM expression levels** | Simulated using beta distributions scaled by circRNA type. | Randomised (using beta distributions) |
| **Mature length** | Computed as sum of selected exon/intron segments. | Computed after simulation |

**Supplementary Table 3.**
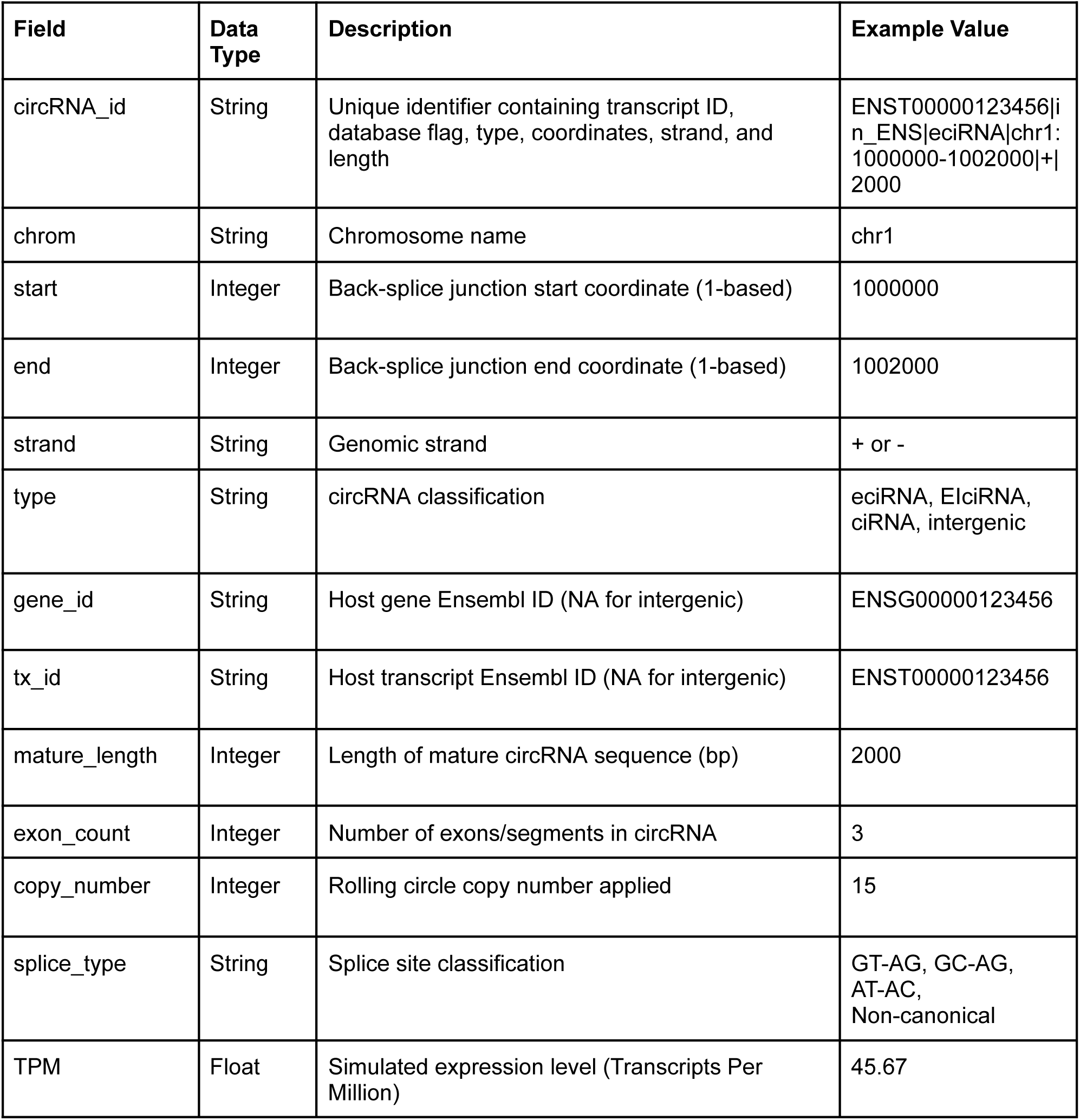
circRNA_metadata.tsv fields used in pipeline for circRNA generation.

**Supplementary Table 4.** Configuration, reference resources, and command-line parameters for circRNA detection tools.

| Parameter | CIRI-long | isoCirc | circNick-LRS |
| --- | --- | --- | --- |
| **Input Requirements** | FASTQ files | FASTQ files | FASTQ files |
| **Reference Genome** | GRCm38.p4.genome_<br>corrected.fa | GRCm38.p4.genome.f<br>a | Built-in:<br>Mus_musculus.GRCm<br>38.87.chr-fix.fa |
| **Version Flexibility** | ✓ User-configurable | ✓ User-configurable | ✗ Fixed (mm10/hg19<br>only) |
| **Gene Annotation** | gencode.vM10.annota<br>tion.gtf | gencode.vM10.annota<br>tion.gtf | Built-in:<br>gencode.vM25.annota<br>tion.gffread.exon.bed |
| **CircRNA Database** | union_bed.bed<br>(circAtlas + circBase) | union_bed.bed<br>(circAtlas + circBase) | Built-in: circAtlas2.0,<br>CIRCpedia_v2,<br>circBase |
| **Database Customization** | ✓ User-configurable | ✓ User-configurable | ✗ Fixed databases<br>only |
| **Provided Threads** | 8 | 8 | 8 (but tool didn't<br>support<br>multithreading) |
| **Command Used** | CIRI-long call<br>-i [reads] -r<br>[ref] -o [out]<br>-t 8 -a [gtf] -c<br>[bed] | python isocirc<br>[reads] [ref]<br>[gtf] [bed]<br>[out] -t 8 | long_read_circRN<br>A run [reads]<br>--species mouse<br>--reference-path<br>[ref]<br>--output-path<br>[out] |

**Supplementary Table 5.**
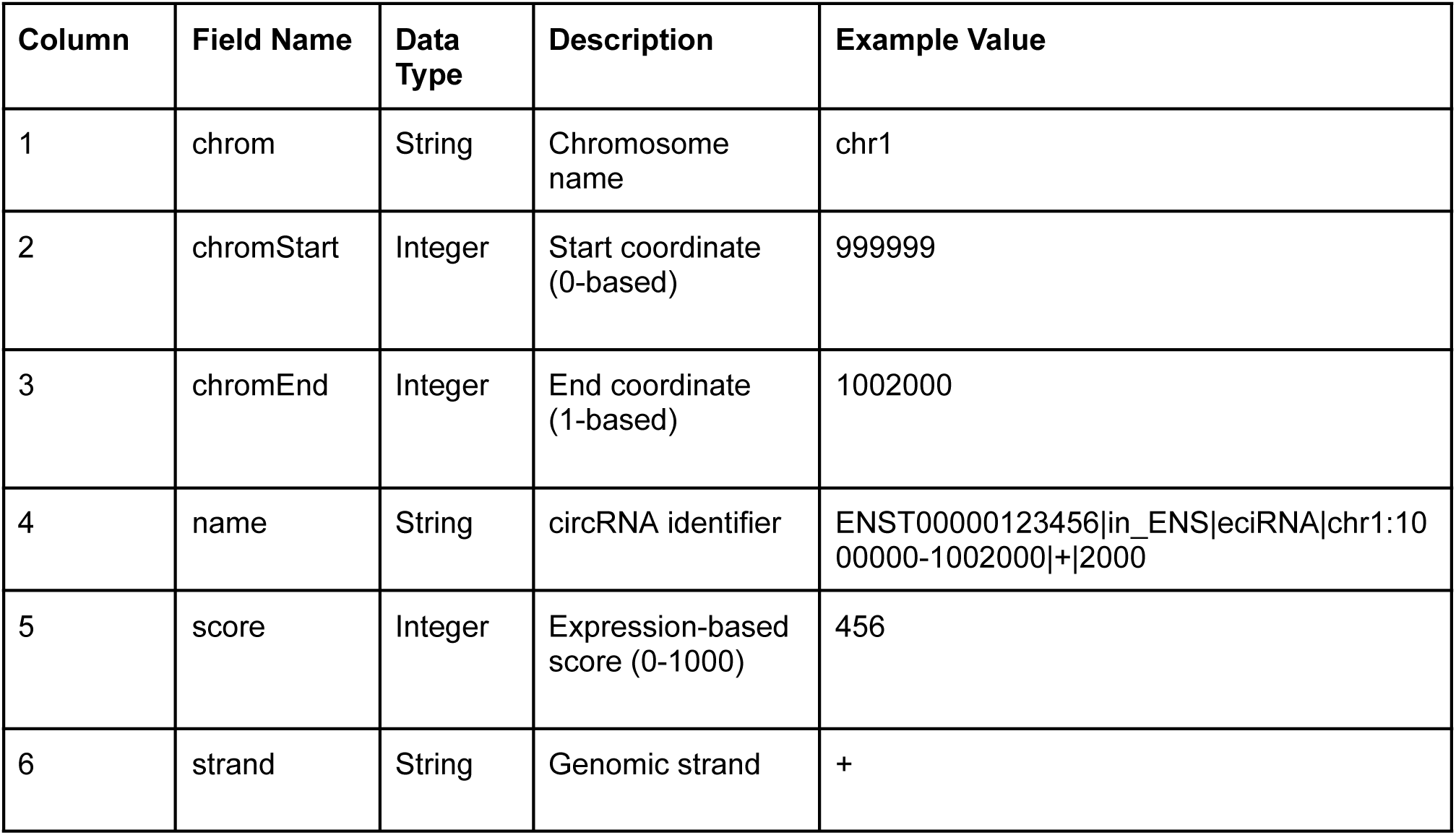
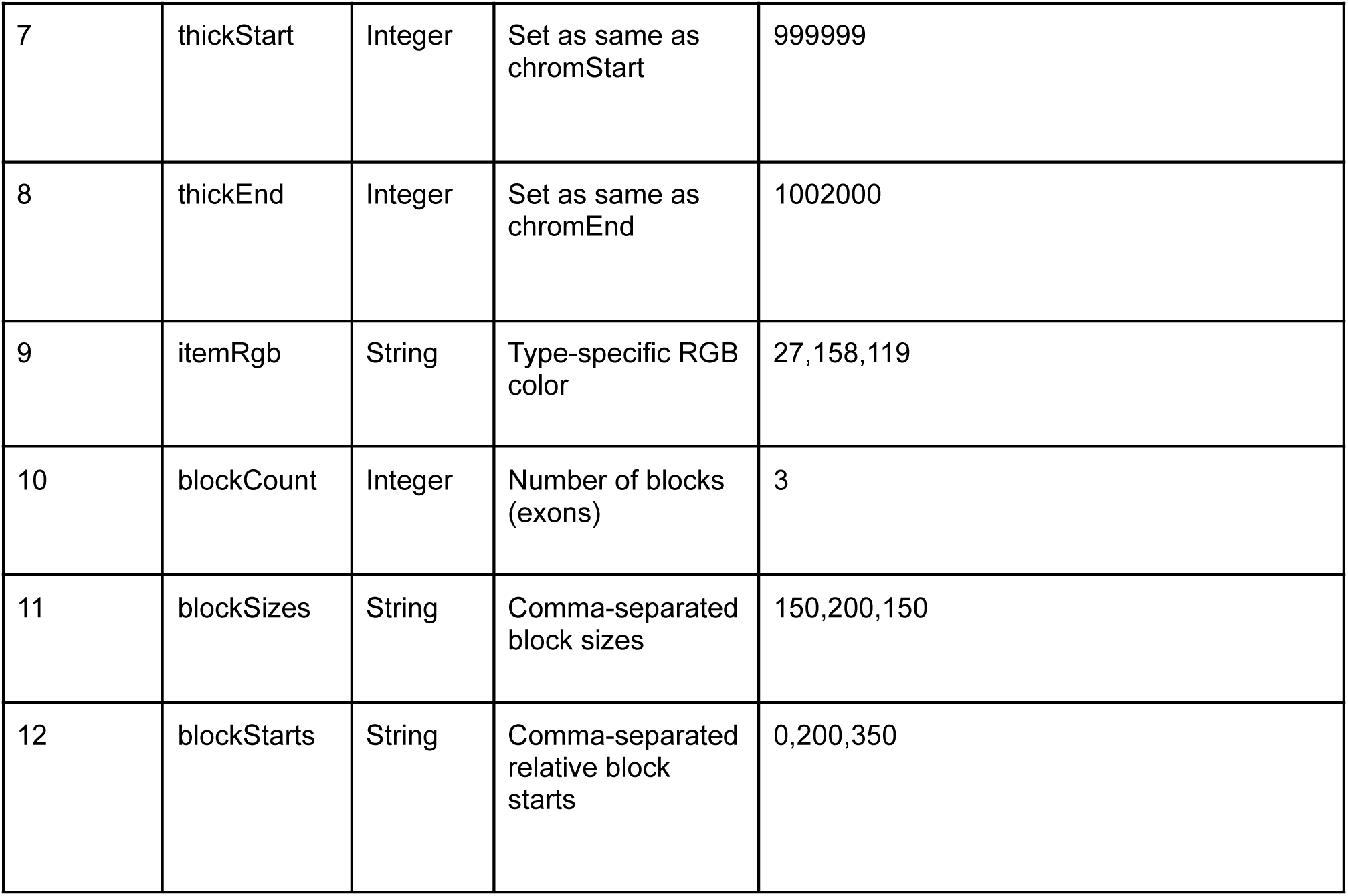
BED12 Format for circRNAs.bed output file used for performance analysis.

## Supplementary materials

### Wet-lab methods

The wet lab methods for these three circRNA detection tools differ significantly in their experimental approaches.

**CIRI-long (Zhang et al., 2021)** uses ribosomal RNA depletion with RiboZero followed by poly(A)-tailing treatment before RNase R digestion for linear RNA removal. The method performs rolling circle reverse transcription with random primers and SMARTer reverse transcriptase to produce long cDNA molecules containing multiple copies of full-length circRNA sequences. Size selection is performed using AMPure XP magnetic beads with different DNA-to-bead ratios (1:1, 1:0.6, and 1:0.5) to capture fragments of approximately 400bp, 600bp, and 1kb respectively, with optimal fragment size selection at approximately 1kb. Libraries are prepared using the ONT Ligation Sequencing Kit (SQK-LSK109).

**isoCirc (Xin et al., 2021)** starts with RNase R treatment to remove linear RNAs, then initiates cDNA synthesis using random hexamers. The circular RNA templates undergo rolling circle amplification (RCA) to generate concatemeric products containing multiple copies of the same circRNA sequence. The RCA products are then debranched using T7 Endonuclease I, subjected to size selection of 3-50kb fragments using BluePippin gel electrophoresis, and libraries are prepared using the ONT Ligation Sequencing Kit (SQK-LSK109).

**circNICK-LRS (Rahimi et al., 2021)** employs ribosomal RNA depletion using RiboZero, followed by RNase R treatment and additional polyadenylation with subsequent poly(A)-depletion to remove remaining linear RNAs. The enriched circRNA pool undergoes controlled fragmentation using NEBNext Magnesium RNA Fragmentation Module at 80°C for variable time periods (30s, 1 min, or 2 min). After fragmentation, 3’ phosphate groups are removed and 5’ phosphate groups are added using T4 Polynucleotide Kinase, followed by polyadenylation. Size selection is performed using agarose gel electrophoresis to isolate PCR products ranging from 350bp to 10kb, and libraries are prepared using the ONT cDNA-PCR Sequencing Kit (SQK-PCS108).

### Tool for ONT in-silico generation of long-read sequencing data

**NanoSim (Hafezqorani et al., 2020):** is a fast and scalable read simulator that captures the technology-specific features of Oxford Nanopore Technologies (ONT) data. The tool analyzes ONT reads from experimental data to model read features such as length, error profiles, and k-mer biases. In its latest version (v3.0), NanoSim supports the simulation of genomic, transcriptomic (cDNA and direct RNA), and metagenomic reads, accommodating features like intron retention events and chimeric reads. The simulation process involves characterizing the input data to learn these features and then generating synthetic reads that mimic the observed characteristics. NanoSim supports the simulation of genomic, transcriptomic (cDNA and direct RNA), and metagenomic reads, accommodating features like intron retention events and chimeric reads. NanoSim is implemented in Python and utilizes tools such as minimap2 for alignment and HTSeq for efficient reading of SAM alignment files. Pre-trained models for organisms like E. coli and S. cerevisiae are available, and users can also train NanoSim on their own datasets to tailor the simulation to specific applications. More details and access to the software can be found at [https://github.com/bcgsc/NanoSim].

### circRNA Databases

**circAtlas v.3 (Wu et al., 2020)**: An integrated resource cataloging over 3 million circRNAs across 33 tissues and 10 vertebrate species. circAtlas 3.0 provides full-length isoform sequences with extensive functional annotations, including conservation profiles, expression patterns, miRNA and RBP binding sites, and coding potential predictions. It integrates both Illumina and Nanopore sequencing data using standardized nomenclature, facilitating cross-database comparisons and evolutionary studies. Its comprehensive tissue and species coverage provides robust reference data for feature extraction and validation.

**circBase (Glažar et al., 2014)**: A foundational database that merges and unifies publicly available datasets of circular RNAs identified in eukaryotic cells. circBase provides comprehensive access to circRNA data within genomic context, supporting queries by identifier, gene, or genomic position. The database includes validation scripts for identifying known and novel circRNAs from sequencing data, making it an essential resource for circRNA research and candidate validation. Its broad taxonomic coverage and condition-independent curation make it ideal for establishing general circRNA feature baselines.

